# MoALE1, an ART-like effector, contributes to rice blast virulence through host immune suppression

**DOI:** 10.64898/2026.11.03.764131

**Authors:** Rachel E. Kalicharan, Nalleli Payne, Isabela G. Tagala, Krzysztof Pawłowski, Jessie Fernandez

## Abstract

Rice blast, caused by the hemibiotrophic fungus *Magnaporthe oryzae*, is one of the most destructive diseases affecting rice production worldwide. During infection, *M. oryzae* secretes effector proteins that suppress host immunity and facilitate colonization. Here, we characterized MoALE1, a predicted effector that is structurally similar to members of the heat-labile enterotoxin/cholera toxin family. Using a yeast secretion trap assay, we demonstrated that MoALE1 contains a functional signal peptide. A MoALE1-GFP fusion localized to the biotrophic interfacial complex, indicating that MoALE1 is a cytoplasmic effector. Expression analyses revealed that MoALE1 is expressed during biotrophy, from 16 to 48 h post-inoculation. Transient expression in *Nicotiana benthamiana* showed that MoALE1 suppresses BAX-mediated cell death, suggesting a role in host immune suppression. To investigate its contribution to pathogenicity, a Δ*moale1* mutant was generated and evaluated in rice infection assays. Deletion of *MoALE1* significantly reduced disease severity but did not affect fungal growth, stress tolerance, sporulation, appressorium development, host penetration, or early invasive growth. In contrast, infection by the Δ*moale1* mutant resulted in enhanced host H₂O₂ accumulation and increased expression of ROS-associated defense genes. Together, these findings identify MoALE1 as a biotrophy-associated cytoplasmic effector that promotes rice blast pathogenicity by suppressing host immune responses and provide new insight into the function of ADP-ribosyltransferase (ART)-like effector candidates in *M. oryzae*.

## 1. Introduction

Fungal phytopathogens pose a detrimental risk to crops, accounting for a loss of over a third of annual production (Almeida et al., 2019; Godfray et al., 2016). One of the most prominent fungal plant pathogens is *Magnaporthe oryzae*, a filamentous ascomycete fungus that infects a variety of grasses, most notably, rice (Fernandez & Orth, 2018; Wilson & Talbot, 2009; Zhang et al., 2022). As the causal agent of rice blast disease, *M. oryzae* wields considerable influence on global rice production, each year destroying up to 30% of harvests (Nalley et al., 2016).

*M. oryzae* follows a hemibiotrophic lifestyle, initially colonizing the plant host in a symptomless manner during its biotrophic phase, before switching to necrotrophy 4-5 days later, resulting in host cell death. To incite pathogenesis, *M. oryzae*’s asexual, three-cell conidia land on the leaf surface and attach via a mucilage substance (Fernandez & Orth, 2018) Natural elements, such as wind or rain, often serve as the vehicle for spore dispersal (Eseola et al., 2021). Following attachment to the leaf, the spore will germinate, forming a germ tube, which eventually differentiates into an infection structure known as the appressorium (Fernandez & Orth, 2018; Osés-Ruiz et al., 2017). The appressorium is melanized, maintaining high turgor pressure, which ultimately facilitates mechanical penetration of *M. oryzae* into host epidermal cells via a penetration peg (De Jong et al., 1997; Kershaw & Talbot, 2009; Veneault-Fourrey et al., 2006). Once inside the host, this peg expands to form a filamentous primary invasive hypha (IH), which later differentiates into a bulbous IH (Kankanala et al., 2007; Khang et al., 2010; Sakulkoo et al., 2018). Furthermore, on this hypha, a membranous structure known as the biotrophic interfacial complex (BIC) will form, which mediates the unconventional secretion of proteinaceous effectors (Giraldo et al., 2013). Effectors secreted from the BIC have been shown to localize to the host cytoplasm and nucleus, altering the host’s immune response to pathogen invasion (Kim et al., 2020; Lee et al., 2023; Park et al., 2012; Park et al., 2016). Additionally, during this time, *M. oryzae* secretes effectors via the conventional ER-Golgi pathway; these effectors localize to the apoplast between the pathogen and host, mediating evasion of host detection (Giraldo et al., 2013; Mentlak et al., 2012; Yang et al., 2019). Together, *M. oryzae*’s repertoire of cytoplasmic and apoplastic effectors plays a crucial role in enabling the pathogen to curb host detection and response, while supporting its proliferation.

As effectors are critical to the pathogen’s ability to establish in the host, circumvent its responses, and ultimately cause disease, they are of prime interest for virulence characterization and potential disease mitigation. Recently, bioinformatic analyses have revealed that *M. oryzae* possesses a subset of effectors predicted to have an ADP-ribosylation fold (Seong & Krasileva, 2021). Proteins with this fold are known as ADP-ribosyltransferases (ART) and catalyze the transfer of ADP-ribose from nicotinamide adenine dinucleotide (NAD^+^) to target molecules such as proteins or nucleic acids (Aravind et al., 2014; Lüscher et al., 2018). In eukaryotes, ADP-ribosylation is a well-documented post-translational modification (PTM) with implications in various vital pathways, including stress response, DNA repair, and cell differentiation (Chappidi et al., 2024; Dasovich & Leung, 2023; Lüscher et al., 2022; Rosado & Pioli, 2021). ADP-ribosylation is also employed by microbial pathogens as a virulence mechanism. Effectors with ADP-ribosyltransferase activity have been extensively characterized in bacterial pathogens, including *Vibrio cholerae, Corynebacterium diphtheriae, Pseudomonas syringae, Legionella pneumophila,* and *Streptomyces scabies*, where they manipulate host cellular processes through ADP-ribosylation of specific targets (Black et al., 2021; Jørgensen et al., 2008; Lyons et al., 2016; Mikolčević et al., 2021; Nicaise et al., 2013; Seong & Krasileva, 2021). ARTC (cholera-toxin-like) and ARTD (diphtheria-toxin-like) catalytic motifs have been documented in these bacterial effectors. To date however, there have been no ART effectors confirmed in fungi.

Of the predicted ART-like effectors in *M. oryzae*, one putative effector, which we designated MoALE1 (<u>A</u>RT-<u>L</u>ike <u>E</u>ffector 1), bears structural similarity to members of the heat-labile enterotoxin/cholera toxin family. Despite the identification of multiple ART-like effector candidates in *M. oryzae*, their biological functions remain largely unknown (Hasnat et al., 2025; Seong & Krasileva, 2023). Here, we characterize the *M. oryzae* effector protein MoALE1. We show that MoALE1 is a biotrophy-associated cytoplasmic effector that localizes to the BIC during infection. Furthermore, we demonstrate that MoALE1 contributes to full pathogenicity in rice blast and suppresses host immune responses. Examination of host responses revealed increased accumulation of host-derived reactive oxygen species (ROS) in Δ*moale1*-infected plants. Collectively, these findings establish MoALE1 as an important virulence-associated effector required for full rice blast pathogenicity and provide a foundation for future investigation of ART-like effectors in fungal pathogens.

## 2. Materials and Methods

### 2.1. Fungal Strains and Growth Conditions

Mutant strains utilized in this study were generated from the wild-type *M. oryzae* strain Guy11 and stored at -20 °C in the laboratory of J. Fernandez at the University of Florida. All fungal strains were grown on complete media (CM) plates agar (50 mL 20x nitrate salts, 1 mL trace elements, 10 g D-glucose, 2 g peptone, 1 g yeast Extract, 1 g casamino acids, 1 mL vitamin solution, pH adjusted to 6.5 with NaOH, 15 g agar) for 7-10 days, at 25 °C and a 12-hour photoperiod. Mutant strains were grown on CM supplemented with at least 100 µg mL⁻¹ of their respective selection.

For stress tolerance assays, 4-mm mycelial plugs were excised from actively growing colonies and transferred to CM supplemented with either 0.5 M NaCl, 1 M sorbitol, 5 mM H₂O₂, 200 μg mL⁻¹ Congo Red, or 0.01% SDS. Plates were incubated for 5 days at 25 °C and imaged using an Epson Workforce scanner at 600 dpi. Colony diameter was measured using Fiji (ImageJ) (Schindelin et al., 2012).

### 2.2. Cloning

The *MoALE1* (MGG_09810) promoter and coding sequence, excluding the stop codon, were amplified from *M. oryzae* Guy11 genomic DNA using the primers listed in Table S1 (Chen et al., 2020; Omar et al., 2016; Spence et al., 2014; Yang et al., 2017). The resulting fragment was assembled in-frame with GFP in the pCB1532 vector using Gibson Assembly (New England Biolabs, Ipswich, MA, USA) following vector linearization with NotI/PstI to generate the pCB1532::MoALE1-GFP construct for localization and complementation studies. For yeast secretion trap assays, full-length *MoALE1* and a signal peptide deletion construct lacking the N-terminal 22 amino acids were amplified and assembled into pYST1 following linearization with EcoRI/NotII. For transient expression assays in *Nicotiana benthamiana*, the *MoALE1* coding sequence was amplified and assembled into the pGD expression vector following digestion with HindIII/PstI. For protein expression and purification via *Escherichia coli*, the *MoALE1* coding sequence was amplified and assembled into the pET28a expression vector following digestion with NotI/SalI. An E143A substitution was introduced into pET28a-*MoALE1* by site-directed mutagenesis using Phusion High-Fidelity DNA Polymerase (Thermo Fisher Scientific). All recombinant plasmids were verified by Sanger sequencing prior to transformation.

### 2.3. Protoplast-mediated Transformation of M. oryzae

Generation of the Δ*moale1* mutant was performed by protoplast-mediated transformation and homologous recombination-mediated gene replacement, as previously described by (Garcia et al., 2023). Briefly, approximately 1-kb regions upstream and downstream of the *MoALE1* coding sequence were amplified by PCR and fused to the *SUR* selectable marker, which confers resistance to sulfonylurea through a mutant *ILV1* allele. The deletion cassette was generated by two-step fusion PCR, in which the left-flank, *SUR* cassette, and right-flank fragments were first amplified independently and subsequently fused into a single construct for transformation.

For protoplast generation, Guy11 mycelia were cultured in liquid CM for 48 h at 25 °C with shaking and digested with *Trichoderma* lysing enzymes in OM buffer (29 g MgSO₄·7H₂O, 1 mL sodium phosphate buffer [pH 5.8], and 0.75 g lysing enzymes from *Trichoderma*; pH adjusted to 5.5 with Na₂HPO₄) to release protoplasts. Protoplasts were purified by sequential washing in ice-cold ST buffer (54.66 g sorbitol and 50 mL 1 M Tris-HCl, pH 7.0) and STC buffer (218.64 g sorbitol, 100 mL 1 M Tris-HCl pH 7.5, and 100 mL 1 M CaCl₂ brought to a final volume of 500 mL with ddH₂O). Purified protoplasts were adjusted to a final concentration of 2-5 × 10⁸ protoplasts mL⁻¹.

For transformation, 100 μL of protoplast suspension was mixed with 1 μg of DNA and 50 μL of STC buffer and incubated at room temperature for 20 min. PEG-mediated transformation was performed by sequential addition of PTC buffer (6 g PEG 4000, 10 μL 1 M Tris-HCl pH 7.5, and 100 μL 1 M CaCl₂ brought to a final volume of 10 mL), followed by a 15-min incubation at room temperature. Transformed protoplasts were suspended in regeneration medium, plated, and incubated overnight prior to overlaying with selective medium containing sulfonylurea. Transformants were recovered at 25 °C and screened by PCR for correct integration of the replacement cassette.

### 2.4. DNA Extractions of M. oryzae Transformants

Mycelial samples from putative *M. oryzae* transformants were collected into 1.5 mL microcentrifuge tubes. Genomic DNA was extracted using a previously described potassium chloride extraction method (Garcia et al., 2023) Briefly, 500 μL of extraction buffer (74.5 g KCl, 100 mL 1 M Tris-HCl [pH 8.0], and 20 mL 0.5 M EDTA [pH 8.0]) was added to each sample, and the fungal tissue was homogenized using a sterile pestle. Samples were centrifuged at 5,000 rpm for 10 min, and the supernatant was transferred to tubes containing 300 μL isopropanol to precipitate DNA. Following centrifugation at 12,000 rpm for 10 min, DNA pellets were washed with 70% ethanol, air-dried, and resuspended in 30 μL nuclease-free water containing 1 μL RNase. Samples were incubated for 15 min at room temperature, followed by 15 min at 55 °C. Genomic DNA was diluted with 250 μL nuclease-free water and used as a template for PCR verification of transformants.

### 2.5. Sporulation Assay

Conidial production was assessed using 10-mm mycelial plugs from Guy11 and Δ*moale1* strains grown on CM for 10 days. Conidia were harvested by flooding plates with sterile water and gently scraping the colony surface. The resulting suspension was filtered through Miracloth, centrifuged, and resuspended in 1 mL sterile water. Conidial concentration was determined using a hemocytometer. Three biological replicates were performed for each strain.

### 2.6. Appressoria Formation Assay

Conidial suspensions were adjusted to a concentration of 1 × 10^5^ conidia mL^-1^ in sterile water. Three 20 μL droplets of conidial suspension were placed onto hydrophobic plastic coverslips and glass coverslips mounted on microscope slides. The slides were incubated in a humid chamber at 24 °C under a 12 h photoperiod. After 16 h, appressorium formation was evaluated microscopically. For each replicate, 50 conidia were scored for the presence or absence of an appressorium. Three biological replicates were performed for each strain.

### 2.7. Pathogenicity and Live-Cell Imaging Assays

Rice plants (*Oryza sativa* cultivars CO39 and YT16) were grown under greenhouse conditions at approximately 25 °C. For spray inoculation assays, conidia were harvested from 10-day-old cultures by flooding plates with sterile water and gently scraping the colony surface. The resulting suspension was filtered through three layers of Miracloth and centrifuged at 3,000 rpm for 5 min. Pelleted conidia were resuspended in sterile water, and spore concentrations were determined using a hemocytometer. Three-and-a-half-week-old rice seedlings were spray-inoculated with conidial suspensions of Guy11 or mutant strains at a final concentration of 5 × 10^4^ conidia mL^-1^ in 0.2% gelatin. Each plant received approximately 5 mL of inoculum. Inoculated plants were maintained at 24 °C under a 12-h photoperiod. Disease symptoms were assessed at 5 dpi by imaging leaves with an Epson Workforce scanner at 600 dpi. Disease severity was quantified as the percentage diseased leaf area using Fiji (ImageJ).

For leaf sheath assays, sheaths from four-week-old rice seedlings were excised and mounted in glass dishes containing moist paper towels to maintain humidity. Sheaths were inoculated with conidial suspensions of Guy11 or the indicated mutant strains at a concentration of 1 × 10^5^ conidia mL^-1^ in 0.2% gelatin. Inoculated sheaths were incubated at 24 °C under a 12-h photoperiod. At the indicated time points, sheaths were cut longitudinally to expose the inner epidermal layer and mounted on slides for microscopic examination.

For DAB staining, infected leaf sheaths were collected at 30 hpi and incubated overnight in 1 mg mL^-1^ 3,3′-diaminobenzidine (DAB) solution pH 3.8 with gentle agitation. Samples were subsequently cleared for 2 h in a 94:4 (w/v) ethanol:acetic acid solution and prepared for microscopy. For DPI treatments, conidial suspensions were treated with 0.4 μM diphenyleneiodonium chloride (DPI) immediately prior to inoculation.

Infected leaf sheaths were visualized with a Zeiss Axio Observer 7 microscope equipped with an Axiocam 503 mono camera. Images were processed using the ZenPro software under the ZEN Tiles and Positions Module (Zeiss).

### 2.8. Suppressor Screen Assay

The BAX-mediated cell death suppression assay was performed as previously described by (Sharpee et al., 2017). Briefly, the pGD expression vector (Goodin et al., 2002), BAX construct, and AvrPiz-t construct were kindly provided by the laboratory of Dr. Guo-Liang Wang. The Mo*ALE1* coding sequence was cloned into the pGD vector, using the primers listed in Table S1, and the resulting constructs were transformed into *Agrobacterium tumefaciens* strain GV3101. A single *A. tumefaciens* colony was cultured in Luria-Bertani (LB) medium (10 g tryptone, 5 g yeast extract, and 5 g NaCl per liter) supplemented with 100 μM acetosyringone (AS) and the appropriate antibiotic selection at 28 °C for 24-48 h. Cultures were adjusted to an OD_600_ of 0.4, pelleted by centrifugation at 3,000 × *g* for 5 min, and washed with infiltration medium (10 mM MgSO₄ and 10 mM MES-KOH, pH 5.6). Cells were resuspended in infiltration medium supplemented with 200 μM AS and incubated at room temperature for at least 3 h prior to infiltration.

Four-week-old *N. benthamiana* leaves were infiltrated with *A. tumefaciens* carrying the MoALE1 expression construct using a needleless syringe. At 24 h post-infiltration, the same leaf areas were challenged with *A. tumefaciens* expressing the mammalian pro-apoptotic protein BAX. Leaf tissue was collected 24 h after BAX infiltration for RNA extraction using the RNeasy Plant Mini Kit (Qiagen), and transgene expression was verified by RT-PCR. At 4 dpi following BAX infiltration, leaves were imaged, and the percentage of cell death at infiltration sites was quantified using Fiji (ImageJ). Buffer-only and GFP controls were included as negative controls, whereas AvrPiz-t served as a positive control for suppression of BAX-mediated cell death.

### 2.9. Yeast Secretion Trap and TTC Assays

The yeast secretion trap assay was performed as previously described by Lee et al. (2006) (S.-J. Lee et al., 2006). Briefly, the full-length (MoALE1_1-239_) and signal peptide-deleted (MoALE1_23-239_) coding sequences were cloned into the pYST expression vector, kindly provided by the laboratory of Dr. Jocelyn K. C. Rose, and transformed into the invertase-deficient Δ*suc2* strain of *Saccharomyces cerevisiae*. Transformants were cultured in SD-Leu glucose medium, adjusted to an OD_600_ of 1.0, and serially diluted. Dilutions were plated onto SD-Leu glucose and SD-Leu sucrose media. Growth on sucrose medium was used to assess secretion of the invertase reporter mediated by the MoALE1 signal peptide. Cultures were incubated at 30 °C for 48 h and imaged using an Epson Workforce scanner at 600 dpi.

Invertase secretion was further evaluated using a 2,3,5-triphenyltetrazolium chloride (TTC) reduction assay as described by (S.-J. Lee et al., 2006). Briefly, yeast transformants were grown in liquid SD-Leu sucrose medium, harvested by centrifugation, washed with sterile water, and incubated in sucrose-containing acetate buffer to promote invertase activity. Culture supernatants were subsequently mixed with TTC solution, and invertase activity was assessed by the formation of a red-colored precipitate resulting from TTC reduction. Development of the red reaction product was interpreted as evidence of functional secretion of the invertase reporter protein.

### 2.10. RNA extraction and qPCR analysis

Total RNA was extracted from infected rice leaf sheath tissues using the RNeasy Plant Mini Kit (Qiagen) according to the manufacturer’s instructions. RNA concentration and quality were assessed spectrophotometrically prior to cDNA synthesis. For each sample, 1 μg of total RNA was reverse transcribed using the qSCRIPT cDNA Synthesis Kit (QuantaBio) following the manufacturer’s protocol.

Quantitative RT-PCR (qPCR) was performed in 10-μL reaction volumes using PerfeCTa SYBR Green FastMix (QuantaBio) and gene-specific primers (Table S1). Reactions were conducted using three biological replicates, each with three technical replicates. Relative transcript abundance was calculated using the comparative Ct (ΔΔCt) method.

For analysis of host gene expression, *OsUBQ10* was used as the endogenous reference gene. For the *MoALE1* infection time-course experiment, the fungal housekeeping gene *MoACT1* was used as the endogenous reference. Relative expression values were calculated relative to the corresponding control treatment.

### 2.11. Protein Purification

The *MoALE1* coding sequence was cloned into the pET28a expression vector using the primers listed in Table S1. Recombinant MoALE1 was expressed in *E. coli* Rosetta cells and induced with 0.4 mM isopropyl β-D-1-thiogalactopyranoside (IPTG) when cultures reached an OD_600_ of 0.6. Following induction, cultures were incubated overnight at 18 °C. Cells were harvested by centrifugation at 5,000 rpm for 20 min at 4 °C and resuspended in lysis buffer (50 mM sodium phosphate, pH 7.4, 300 mM NaCl, 10 mM imidazole, 0.5% Triton X-100, 1× β-mercaptoethanol, 25 mg mL^−1^ lysozyme, 1 mM PMSF, and Benzonase). Following incubation on ice for 20 min, cells were lysed by sonication and clarified by centrifugation at 15,000 rpm for 20 min at 4 °C.

Recombinant His-tagged MoALE1 was purified using Ni-NTA agarose resin (Thermo Fisher Scientific). Bound protein was washed with wash buffer (50 mM sodium phosphate, pH 7.4, 300 mM NaCl, 10 mM imidazole) and eluted with elution buffer (50 mM sodium phosphate, pH 7.4, 300 mM NaCl, 250 mM imidazole). Eluted fractions were concentrated and further purified by size-exclusion chromatography using a Superdex 200 column (Cytiva) equilibrated in 50 mM sodium phosphate, pH 7.4, 300 mM NaCl, 5% glycerol, and 1 mM DTT. Peak fractions were pooled, aliquoted, and stored at −80 °C. Protein purity was assessed by SDS-PAGE.

### 2.12. Immunoblot Analysis

Proteins were separated by 10% SDS-PAGE and transferred to polyvinylidene difluoride (PVDF) membranes. MoALE1 expression was detected using an HRP-conjugated anti-His antibody (Proteintech; 1:10,000 dilution) in 5% milk overnight. Membranes were washed three times with 1× TBST and subsequently incubated with ProSignal Pico chemiluminescent substrate (Genesee Scientific). Chemiluminescent signals were visualized using an iBright FL1500 imaging system (Thermo Fisher Scientific).

### 2.13. Bioinformatic Analyses

FFAS and HHpred were used to identify similarity of MoALE1 to ADP-ribosyltransferases (Xu et al., 2014; Zimmermann et al., 2018). SignalP version 6.0 was used to predict the presence and cleavage site of the MoALE1 signal peptide (Teufel et al., 2022). Multiple sequence alignments were generated using Clustal Omega and visualized with ESPript 3.0 (Robert et al., 2025; Sievers et al., 2011). Structural prediction of MoALE1 and molecular docking analyses with NAD⁺ were performed using AlphaFold 3. Predicted structures and docking models were visualized, analyzed, and rendered using UCSF ChimeraX (Abramson et al., 2024; Meng et al., 2023).

### 2.14. Statistical Analysis

Statistical analyses were performed using GraphPad Prism (GraphPad Software, San Diego, CA, USA). All experiments were conducted using at least three independent biological replicates unless otherwise stated, and data are presented as the mean ± standard deviation (SD). Comparisons between two groups were analyzed using a two-tailed Student’s *t*-test, whereas comparisons among multiple groups were analyzed using one-way analysis of variance (ANOVA) followed by Dunnett’s multiple-comparison test using the appropriate control group. Differences were considered statistically significant at *P* < 0.05. Statistical significance is indicated in figures as follows: ns, not significant (*P* ≥ 0.05); * (*P* < 0.05); ** (*P* < 0.01); *** (*P* < 0.001); and **** (*P* < 0.0001). Graphs were generated using GraphPad Prism, and schematic illustrations were created using BioRender.com.

## 3. Results

### 3.1. *M. oryzae* Encodes an ARTC-Like Protein

Initial sequence analysis of MoALE1 (MGG_09810) by FFAS and HHpred revealed that it is a 239-amino-acid protein containing a predicted N-terminal signal peptide and an Arg-Ser-Glu-x-Glu (R-S-ExE) motif characteristic of mono-ADP-ribosyltransferases (mARTs) (Fig. 1A). To further investigate its relationship to known mARTs, MoALE1 was compared with the catalytic A subunits of cholera toxin (*Vibrio cholerae* CtxA) and heat-labile enterotoxin (*E. coli* EltA). Sequence alignment revealed conservation of residues corresponding to the R-S-ExE motif and other regions associated with the catalytic core of these toxins, suggesting structural similarity between MoALE1 and members of the heat-labile enterotoxin/cholera toxin family (Fig. 1B). Recombinant expression of MoALE1 in *E. coli* yielded a His-tagged protein of the expected molecular weight (∼29 kDa), as confirmed by SDS-PAGE and immunoblot analysis (Fig. S1).

**Figure 1.**
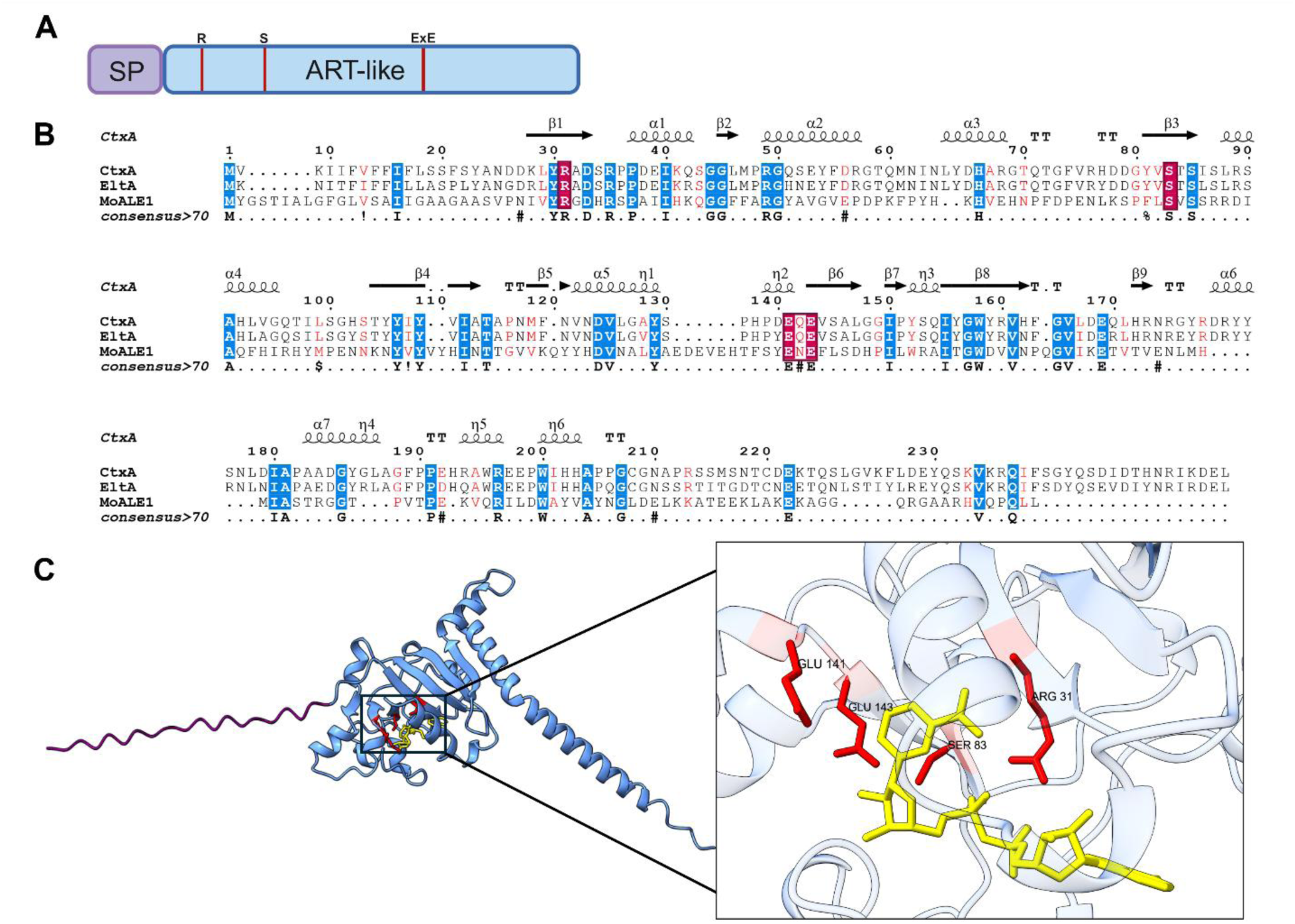
Structural analysis of MoALE1 reveals an ART-like domain with similarity to bacterial enterotoxins. **(A)** Schematic representation of MoALE1 showing the predicted N-terminal signal peptide (SP) and ART-like domain. Conserved residues corresponding to the R-S-ExE motif are indicated. Created using BioRender.com. **(B)** Multiple sequence alignment of MoALE1 with the catalytic A subunits of cholera toxin (CtxA) and heat-labile enterotoxin (EltA). Conserved residues are highlighted, and residues comprising the predicted R-S-ExE motif are boxed in red. Secondary structure elements are shown above the alignment. **(C)** AlphaFold3 structural model of MoALE1. The signal peptide is shown in purple, and the ART-like domain is shown in blue. Predicted NAD⁺ is shown in yellow. Inset: enlarged view of the putative NAD⁺-binding pocket. Conserved residues R31, S83, E141, and E143 are highlighted in red and shown in proximity to the predicted NAD⁺ molecule.

To evaluate whether the predicted ART-like fold of MoALE1 could accommodate NAD⁺, molecular docking was performed using AlphaFold-generated models. The predicted MoALE1-NAD⁺ complex yielded an ipTM score of 0.95. Structural analysis indicated that NAD⁺ was positioned within the predicted ligand-binding pocket, with residues of the conserved R-S-ExE motif located in close proximity to the molecule (Fig. 1C). Structural superposition of MoALE1 with CtxA and EltA revealed strong structural conservation, with RMSD values of 0.891 Å and 1.086 Å, respectively (Fig. S2). The overall fold and spatial arrangement of residues surrounding the predicted NAD⁺-binding pocket were highly conserved among the three proteins. Collectively, these analyses indicate that MoALE1 possesses structural features consistent with an ART-like fold and shares similarity with members of the heat-labile enterotoxin/cholera toxin family.

### 3.2. *MoALE1* Encodes a Functional Signal Peptide

Following the bioinformatic assessment of MoALE1, we began functional characterization of the effector. The MoALE1 sequence was analyzed using SignalP 6.0, which predicted a 94% probability of a signal peptide comprising the first 22 amino acids (Teufel et al., 2022) (Fig. S3). To experimentally assess signal peptide function, we performed a yeast secretion trap assay (Lee & Rose, 2012). The full-length sequence of MoALE1 containing its N-terminal signal peptide (MoALE1_1-239_) and a truncated variant lacking the signal peptide (MoALE1_23-239_) were each fused to a truncated invertase (*suc2*) reporter lacking its native N-terminal signal peptide. These constructs were transformed into the *S. cerevisiae* BY4741 *Δsuc2* mutant strain.

Transformants were plated on SD-Leu glucose, which served as a growth control, and SD-Leu sucrose, which required secretion of functional invertase for growth. Wild-type BY4741 carrying the empty vector grew on both media, whereas the Δ*suc2* strain carrying the empty vector grew only on SD-Leu glucose. The *M. oryzae* effector Bas1 was included as a positive control; Bas1_1-115_ grew on both media, whereas Bas1_23-115_ failed to grow on SD-Leu sucrose. Similarly, yeast expressing MoALE1_1-239_ grew on both media, whereas MoALE1_23-239_ failed to grow on SD-Leu sucrose (Fig. 2A).

**Figure 2.**
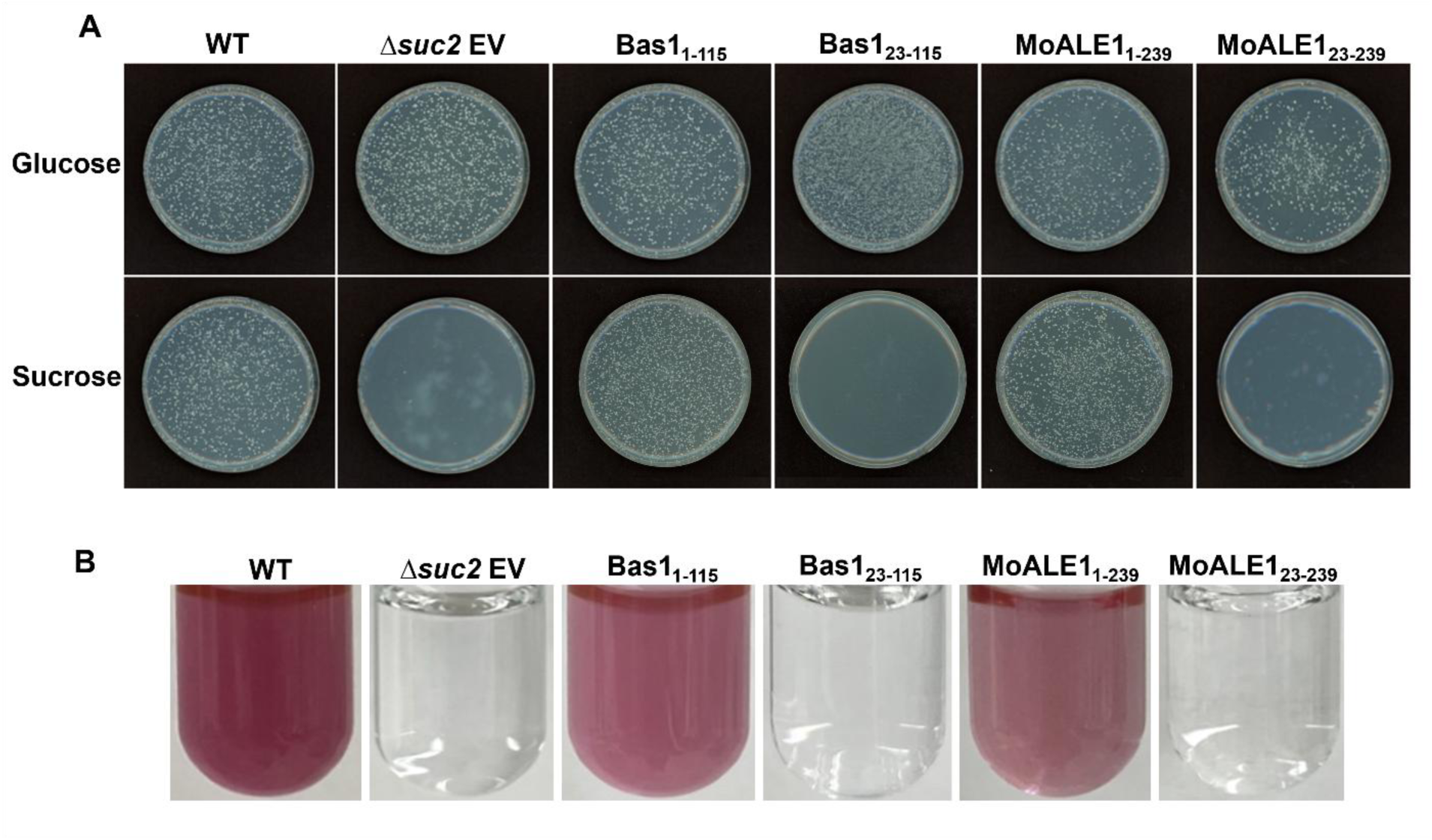
MoALE1 contains a functional signal peptide. **(A)** Yeast secretion trap assay using the invertase-deficient Δ*suc2* strain of *Saccharomyces cerevisiae*. Transformants were grown on SD-Leu glucose and SD-Leu sucrose media to assess invertase reporter secretion. WT (BY4741) and Δ*suc2* carrying the empty vector (EV) served as positive and negative controls, respectively. Bas1 was included as a positive control for secretion. Constructs containing the native signal peptide (Bas1_1-115_ and MoALE1_1-239_) supported growth on sucrose medium, whereas constructs lacking the signal peptide (Bas1_23-115_ and MoALE1_23-239_) failed to grow. Representative plates from the 1:100 dilution are shown. **(B)** 2,3,5-Triphenyltetrazolium chloride (TTC) reduction assay used to confirm invertase secretion shown in (A). Red coloration indicates secretion of invertase into the culture medium, whereas the absence of color development indicates a lack of invertase secretion.

To further validate the yeast secretion trap results, a TTC assay was performed. In this assay, secreted invertase reduces colorless 2,3,5-triphenyltetrazolium chloride (TTC) to a red, insoluble product. Consistent with the growth assays, wild-type BY4741 empty vector, and both full-length Bas1 and MoALE1 exhibited a strong colorimetric change, whereas strains lacking a functional signal peptide did not (Fig. 2B). Together, these results demonstrate that MoALE1 contains a functional signal peptide that mediates protein secretion.

### 3.3. MoALE1 Is Expressed During Biotrophy and Localizes to the BIC

To examine *MoALE1* expression and localization during infection, a *MoALE1*-GFP fusion driven by the native *MoALE1* promoter was generated and introduced into *M. oryzae*. Fluorescence was not detected in vegetative hyphae, conidia, or appressoria, indicating that *MoALE1* is not expressed during vegetative growth or appressorium development (Fig. 3A).

**Figure 3.**
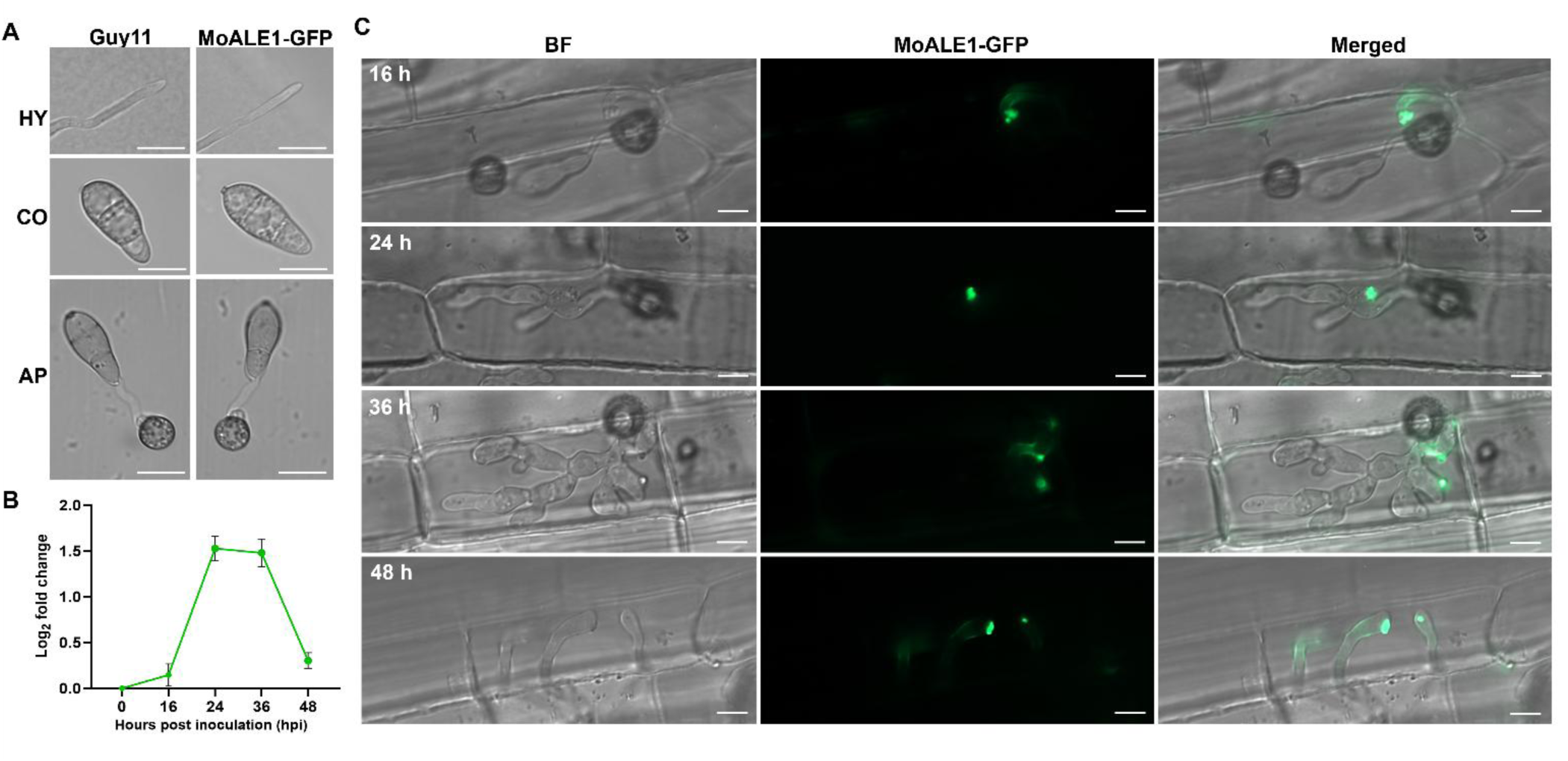
MoALE1 is expressed during biotrophic growth and localizes to the BIC. **(A)** Merged bright-field and fluorescent channel images of Guy11 and the MoALE1-GFP strain during vegetative hyphal growth (HY), conidial development (CO), and appressorium formation (AP). No GFP fluorescence was detected in vegetative hyphae, conidia, or appressoria. **(B)** Relative *MoALE1* transcript abundance during rice infection determined by RT-qPCR. Expression levels are shown as log₂ fold change relative to 0 hpi following normalization to *MoACT1*. Data represent mean ± SD of three biological replicates. **(C)** Subcellular localization of MoALE1-GFP during rice sheath infection at 16, 24, 36, and 48 hpi. Bright-field (BF), GFP fluorescence, and merged images are shown. White arrows indicate BIC-associated MoALE1-GFP fluorescence. Fluorescent signals not indicated by arrows in the 48 hpi panel represent background fluorescence. Scale bars = 10 μm.

To determine the temporal expression pattern of *MoALE1* during infection, transcript abundance was quantified by RT-qPCR at 0, 16, 24, 36, and 48 hpi. *MoALE1* expression increased during infection, peaking at 24 and 36 hpi, then declining at 48 hpi; however, transcript abundance remained higher than at 0 hpi (Fig. 3B). These results indicate that *MoALE1* is preferentially expressed during the biotrophic stage of infection.

We next examined the subcellular localization of MoALE1 during host colonization. Rice leaf sheaths inoculated with the MoALE1-GFP strain were observed at 16, 24, 36, and 48 hpi. Fluorescence was first detected during early infection and consistently accumulated at the biotrophic interfacial complex (BIC) (Fig. 3C). At 24 hpi, MoALE1-GFP localized to BIC-associated invasive hyphae within the first infected cell. At later time points, fluorescence remained associated with BICs located at the tips of invasive hyphae as they extended into neighboring host cells, indicating continued localization during cell-to-cell colonization. Although fluorescence was still detectable at 48 hpi, signal intensity was reduced relative to earlier time points, consistent with the expression profile observed by RT-qPCR (Fig. 3B, C). Together, these results demonstrate that MoALE1 is a biotrophy-associated cytoplasmic effector that is expressed during early infection and accumulates at the BIC throughout biotrophic growth.

### 3.4. MoALE1 Suppresses BAX-Mediated Cell Death

As numerous *M. oryzae* effectors suppress host cell death to promote infection, we investigated whether MoALE1 could inhibit BAX-mediated cell death in *Nicotiana benthamiana*. MoALE1 was transiently expressed in *N. benthamiana* via agroinfiltration, with buffer and GFP serving as negative controls and the *M. oryzae* effector AvrPiz-t as a positive control (Zhong et al., 2018). At 24 h post-infiltration, leaves were challenged with the cell death inducer BAX at the same infiltration sites (Sharpee et al., 2017). 24 h after BAX infiltration, leaf tissue was harvested to verify transgene expression by RT-PCR (Fig. 4A). Leaves were subsequently imaged at 4 dpi, and the percentage of cell death at infiltration sites was quantified (Fig. 4B, C). As expected, buffer- and GFP-treated samples exhibited extensive BAX-induced cell death, averaging approximately 89% and 81%, respectively (Fig. 4B, C). In contrast, MoALE1 expression reduced BAX-induced cell death to nearly undetectable levels (∼1%), comparable to the suppression observed with the positive control AvrPiz-t (∼1%). No significant difference was detected between MoALE1 and AvrPiz-t, whereas both treatments exhibited significantly lower levels of cell death than the buffer and GFP controls (Fig. 4B, C). These findings demonstrate that MoALE1 effectively suppresses BAX-mediated cell death and suggest that it functions as an immune-suppressing effector, attenuating host defense-associated cell death responses.

**Figure 4.**
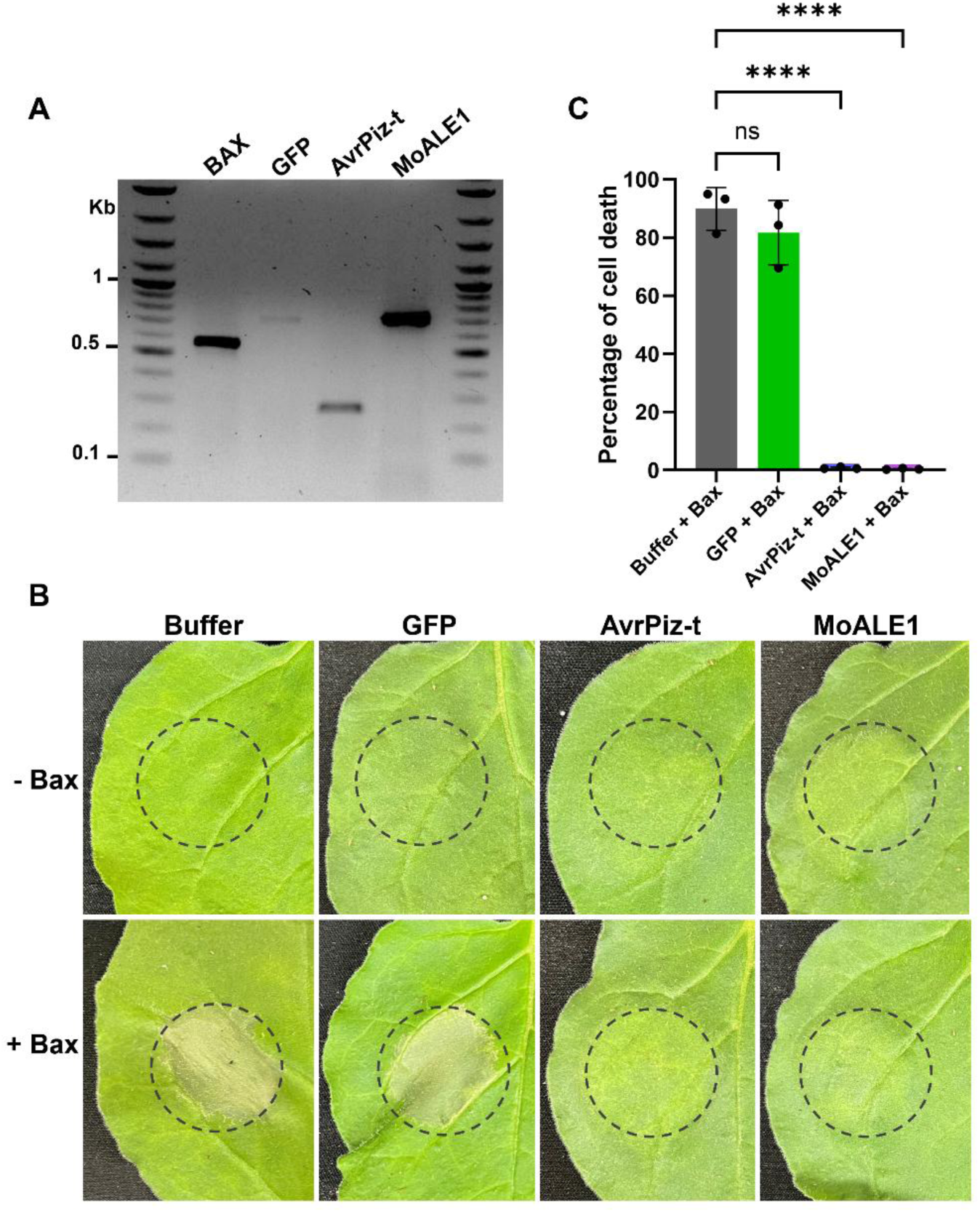
MoALE1 suppresses BAX-mediated cell death. **(A)** RT-PCR confirmation of transgene expression in infiltrated *Nicotiana benthamiana* leaves. Expression of *BAX*, *GFP*, *AvrPiz-t*, and *MoALE1* was verified following agroinfiltration. **(B)** BAX suppression assay in *N. benthamiana*. Four-week-old leaves were infiltrated with buffer, GFP (negative control), AvrPiz-t (positive control), or MoALE1. Twenty-four hours later, the same infiltration sites were challenged with *Agrobacterium tumefaciens* expressing the mammalian cell death inducer BAX. Images were captured 4 dpi following BAX infiltration. Dashed circles indicate infiltration sites. **(C)** Quantification of cell death at infiltration sites. The percentage of cell death was determined from infiltrated leaf areas at 4 dpi. Buffer- and GFP-treated leaves exhibited extensive BAX-induced cell death, whereas expression of MoALE1 or AvrPiz-t significantly reduced cell death. Each dot represents an independent biological replicate (*n* = 3). Bars represent the mean ± SD. Statistical significance was determined using a one-way ANOVA followed by Dunnett’s multiple-comparisons test (ns, not significant; ****, *P* < 0.0001).

### 3.5. *MoALE1* contributes to Full Rice Blast Pathogenicity

To evaluate the contribution of MoALE1 to *M. oryzae* virulence, Δ*moale1* mutants were generated in the wild-type strain Guy11 by replacing the endogenous gene with a selectable marker (Fig. S4). Following PCR confirmation of the deletion mutants and generation of complemented strains, pathogenicity assays were performed on the moderately resistant rice cultivar CO39 and the susceptible rice cultivar YT16. Disease severity was assessed five days after spray inoculation.

In the CO39 cultivar, Guy11-inoculated plants exhibited extensive disease symptoms, with lesions covering approximately 43% of the leaf area (Fig. 5A, B). In contrast, plants inoculated with Δ*moale1* displayed a significant reduction in disease severity, with lesions covering only ∼16% of the leaf area. Complementation of the mutant restored pathogenicity to near wild-type levels (∼39%), confirming that the reduced virulence phenotype resulted from loss of *MoALE1*. A similar trend was observed in the susceptible cultivar YT16 (Fig. 5C, D). Plants inoculated with Guy11 exhibited approximately 28% disease coverage, whereas Δ*moale1*-inoculated plants displayed substantially reduced disease symptoms (∼7%). Restoration of *moale1* in the complementation strain recovered disease severity to approximately 28%, comparable to that observed for the wild-type strain (Fig. 5D). The reduction in disease severity observed in both rice cultivars, together with restoration of the phenotype in the complemented strain, demonstrates that MoALE1 contributes to full virulence of *M. oryzae* during rice infection.

**Figure 5.**
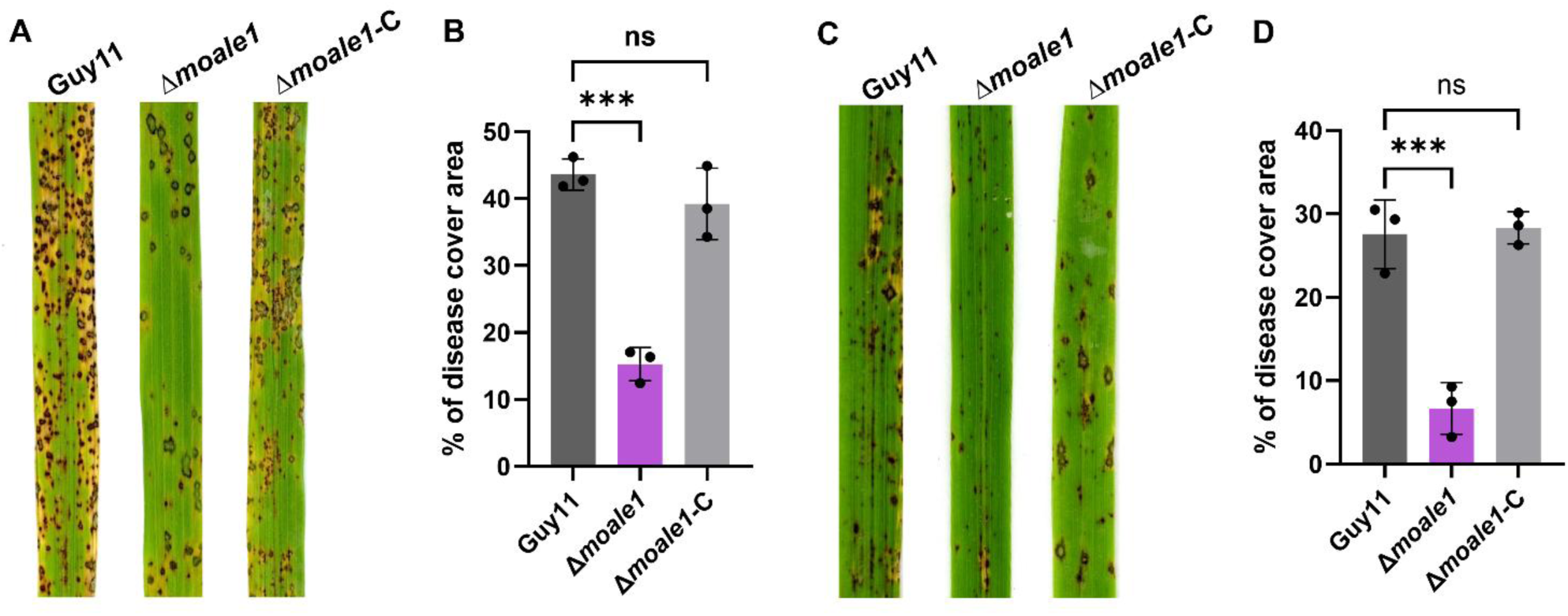
MoALE1 contributes to full rice blast virulence. **(A, C)** Spray inoculation assays on 3.5-week-old rice cultivars CO39 and YT16, respectively, with Guy11 (wild type), Δ*moale1* (deletion mutant), and Δ*moale1*-C (complemented strain). Disease symptoms were assessed at 5 dpi. **(B, D)** Quantification of disease severity shown in (A) and (C), respectively, presented as a percentage of disease-covered leaf area. Each point represents an independent biological replicate (*n* = 3). Bars represent the mean ± SD. Statistical significance was determined using a one-way ANOVA followed by Dunnett’s multiple-comparisons test (ns, not significant; ***, *P* < 0.001).

### 3.6. *MoALE1* Is Dispensable for Fungal Growth and Early in planta Development

Following the observation that deletion of *MoALE1* reduced rice blast disease severity, we sought to determine whether this phenotype resulted from defects in fungal growth, development, or infectivity. Vegetative growth of Guy11 and Δ*moale1* mutants was first assessed on complete medium and under a range of stress conditions, including osmotic stress (NaCl and sorbitol), oxidative stress (H₂O₂), and cell wall stress (SDS and Congo Red). No differences in colony morphology or growth were observed among the strains under any condition tested, indicating that *MoALE1* is dispensable for vegetative growth and stress tolerance (Fig. S5A, B).

To determine whether *MoALE1* influences asexual development, sporulation rates were quantified and were comparable between Guy11 and the Δ*moale1* mutants (Fig. S5C). We further examined appressorium development and morphology and found no significant differences among strains, indicating that deletion of *MoALE1* does not impair appressorium formation (Fig. S5D, E).

Because appressorium development appeared normal, we next assessed infection progression during the early stages of host colonization. Appressorium penetration rates in rice leaf sheaths were comparable among Guy11 and the Δ*moale1* mutants, demonstrating that loss of *MoALE1* does not affect host penetration (Fig. S5F, G). We then quantified invasive hyphal development at 24 hpi and observed no significant differences in the progression of infection stages among strains (Fig. S5H). Similarly, by 48 hpi, invasive hyphae of the Δ*moale1* mutants spread into neighboring cells at frequencies comparable to those of the wild-type strain, indicating that cell-to-cell movement was unaffected by deletion of *MoALE1* (Fig. S5I). These findings indicate that MoALE1 is dispensable for fungal growth, development, and early in planta infection. Therefore, the reduced virulence observed in the Δ*moale1* mutant is unlikely to result from defects in fungal fitness or early infection processes.

### 3.7. *MoALE1* Modulates Host ROS Generation During Infection

Because deletion of *MoALE1* did not affect fungal growth, development, or early infection progression, we next investigated whether the reduced virulence of the Δ*moale1* mutant was associated with altered host defense responses. To assess ROS accumulation, rice leaf sheaths infected with Guy11 or Δ*moale1* strains were subjected to 3,3′-diaminobenzidine (DAB) staining at 30 hpi. Cells infected with Δ*moale1* exhibited markedly stronger DAB staining and increased brown precipitate accumulation relative to Guy11-infected cells, indicating elevated H₂O₂ production (Fig. 6A). Treatment of spores with the NADPH oxidase inhibitor diphenyleneiodonium chloride (DPI) abolished DAB staining in both wild-type and mutant infections, confirming that the observed ROS accumulation was host-derived.

**Figure 6:**
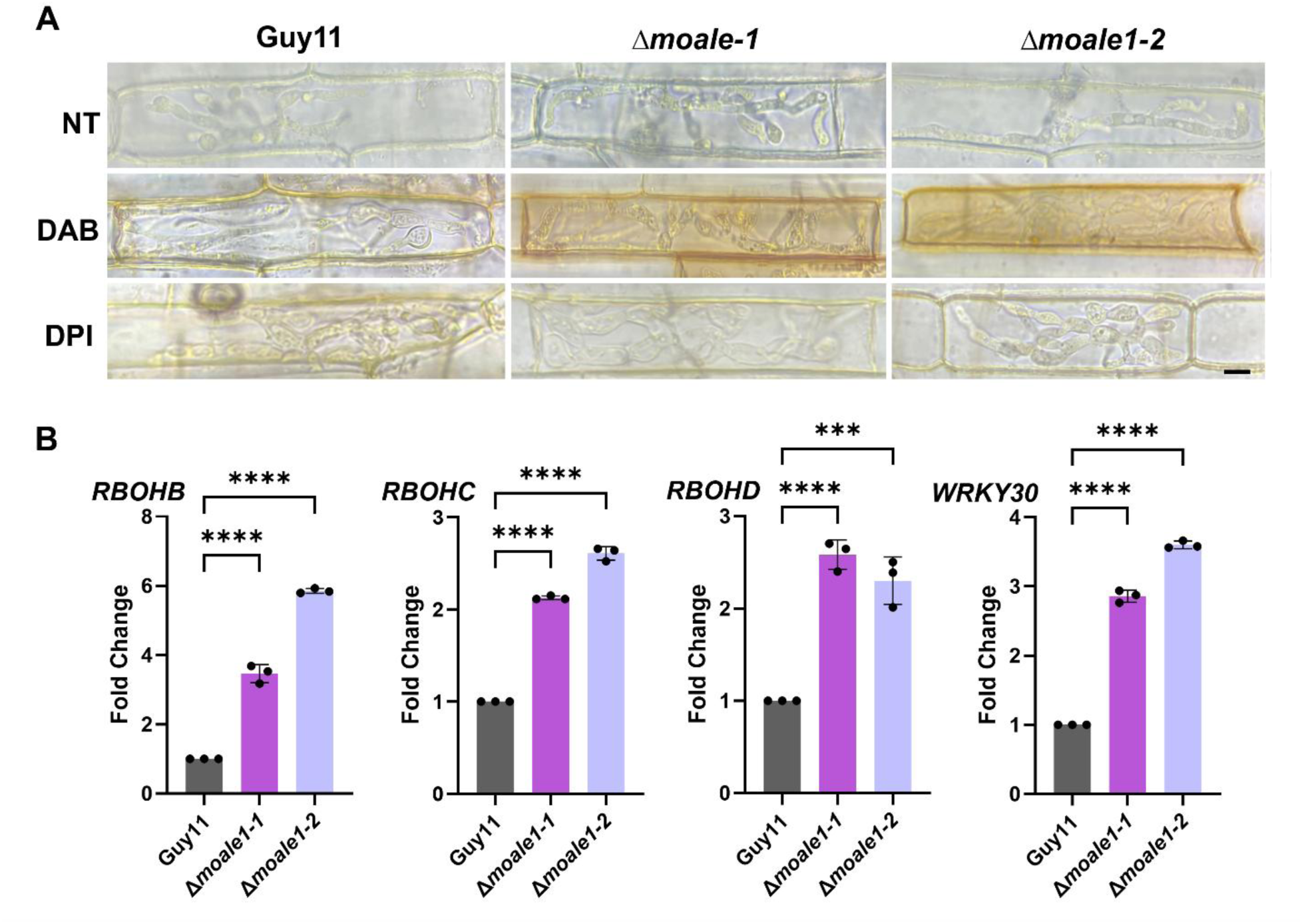
*MoALE1* suppresses host-derived ROS. **(A)** 3.5-week-old YT16 rice leaf sheaths were infected with Guy11 or Δ*moale1-1* and Δ*moale1-2* split marker deletion strains and stained with 3,3’-diaminobenzidine (DAB) 30hpi to visualize host-generated H_2_O_2_. Brown precipitate indicates accumulation of H_2_O_2_ compared to unstained no treatment (NT) and stained diphenyleneiodonium (DPI) controls. Scale bar = 5µm. **(B)** qPCR analysis of relative expression of host NADPH oxidase genes *RBOHB*, *RBOHC*, *RBOHD* and defense gene *WRKY30* between Guy11 and Δ*moale1*-infected leaves (5 dpi). Statistical significance was determined using a one-way ANOVA followed by Dunnett’s multiple-comparisons test (*** indicates *P* < 0.001, **** indicates *P* < 0.0001). Each dot represents an independent biological replicate (*n* = 3), calculated as the mean of three technical replicates. Bars indicate mean ± SD.

To further determine whether enhanced ROS accumulation was associated with activation of ROS-related defense pathways, expression of several host defense genes was analyzed by qPCR. Compared with Guy11-infected plants, tissues infected with the Δ*moale1* mutants exhibited significantly increased expression of the NADPH oxidases *RBOHB*, *RBOHC*, and *RBOHD* (Fig. 6B). These genes contribute to ROS generation and accumulation during immune responses (Khaskhali et al., 2025; Peng et al., 2012).In addition, *WRKY30* expression, a positive regulator of ROS-mediated defense signaling, was significantly elevated in Δ*moale1*-infected tissues (Fig. 6B). The consistent upregulation of these ROS-associated genes further supports the enhanced activation of host defense responses in the absence of MoALE1. Together, the DAB staining and gene expression analyses indicate that the loss of *MoALE1* leads to increased host H₂O₂ accumulation and activation of ROS-associated defense pathways during infection. These findings suggest that *MoALE1* contributes to *M. oryzae* virulence by suppressing H₂O₂-mediated host immune responses.

## 4. Discussion

Rice blast disease, caused by *M. oryzae*, remains one of the most destructive diseases affecting rice production worldwide. Although *M. oryzae* is predicted to secrete hundreds of effectors during infection, the functions of many remain poorly understood. Here, we provide experimental characterization of MoALE1 (ART-Like Effector 1), a previously uncharacterized *M. oryzae* effector possessing structural features consistent with an ADP-ribosyltransferase-like fold. While its biochemical activity remains unknown, our findings establish MoALE1 as a biotrophy-associated cytoplasmic effector that contributes to rice blast pathogenicity and suppresses host immune responses.

Several observations support the classification of MoALE1 as a cytoplasmic effector. We demonstrated that MoALE1 contains a functional signal peptide capable of mediating secretion and that the protein localizes to the BIC during infection. Furthermore, expression analyses revealed that MoALE1 is specifically expressed during biotrophy, with transcript accumulation and fluorescence detected from 16 to 48 hpi. Localization of MoALE1 to the BIC throughout this period strongly supports its classification as a cytoplasmic effector and suggests its involvement in the early stages of host colonization (Giraldo et al., 2013; Kim et al., 2020; Lee et al., 2023; Park et al., 2012; Park et al., 2016). Similar expression and localization patterns have been reported for several *M. oryzae* cytoplasmic effectors, including Bas1 and Mep2, which accumulate at the BIC during biotrophic growth (Mosquera et al., 2009; Yan et al., 2023). Although the biological functions of many BIC-localized effectors remain unresolved, several characterized cytoplasmic effectors have been shown to manipulate host immunity by targeting immune signaling pathways, transcriptional regulation, and cell death responses. Consistent with this model, MoALE1 suppressed BAX-mediated cell death in tobacco leaves, supporting a role for MoALE1 in suppression of host immune responses during biotrophic colonization.

Pathogenicity assays further established MoALE1 as an important virulence-associated effector. Deletion of *MoALE1* significantly reduced disease severity in both the moderately resistant cultivar CO39 and the susceptible cultivar YT16, while complementation restored virulence. Importantly, the reduced pathogenicity of the Δ*moale1* mutant could not be attributed to defects in fungal growth or development. Vegetative growth, stress tolerance, sporulation, appressorium formation, penetration efficiency, invasive hyphal growth, and cell-to-cell spread were all comparable between wild-type and mutant strains. These observations indicate that *MoALE1* contributes to pathogenicity by modulating host responses rather than by affecting fungal fitness or infectivity.

One of the most striking phenotypes associated with loss of *MoALE1* was enhanced host H₂O₂ accumulation during infection. DAB staining revealed substantially stronger H₂O₂ accumulation in tissues infected with the Δ*moale1* mutant relative to the wild-type strain. Moreover, treatment with the NADPH oxidase inhibitor DPI abolished this phenotype, confirming that the observed ROS originated from host cells. Consistent with these observations, plants infected with the Δ*moale1* mutant exhibited elevated expression of the NADPH oxidases *RBOHB*, *RBOHC*, and *RBOHD*, as well as the defense-associated transcription factor *WRKY30*. Together, these findings indicate that MoALE1 suppresses host H₂O₂ accumulation and ROS-associated defense responses during infection (Fig. 7).

**Figure 7.**
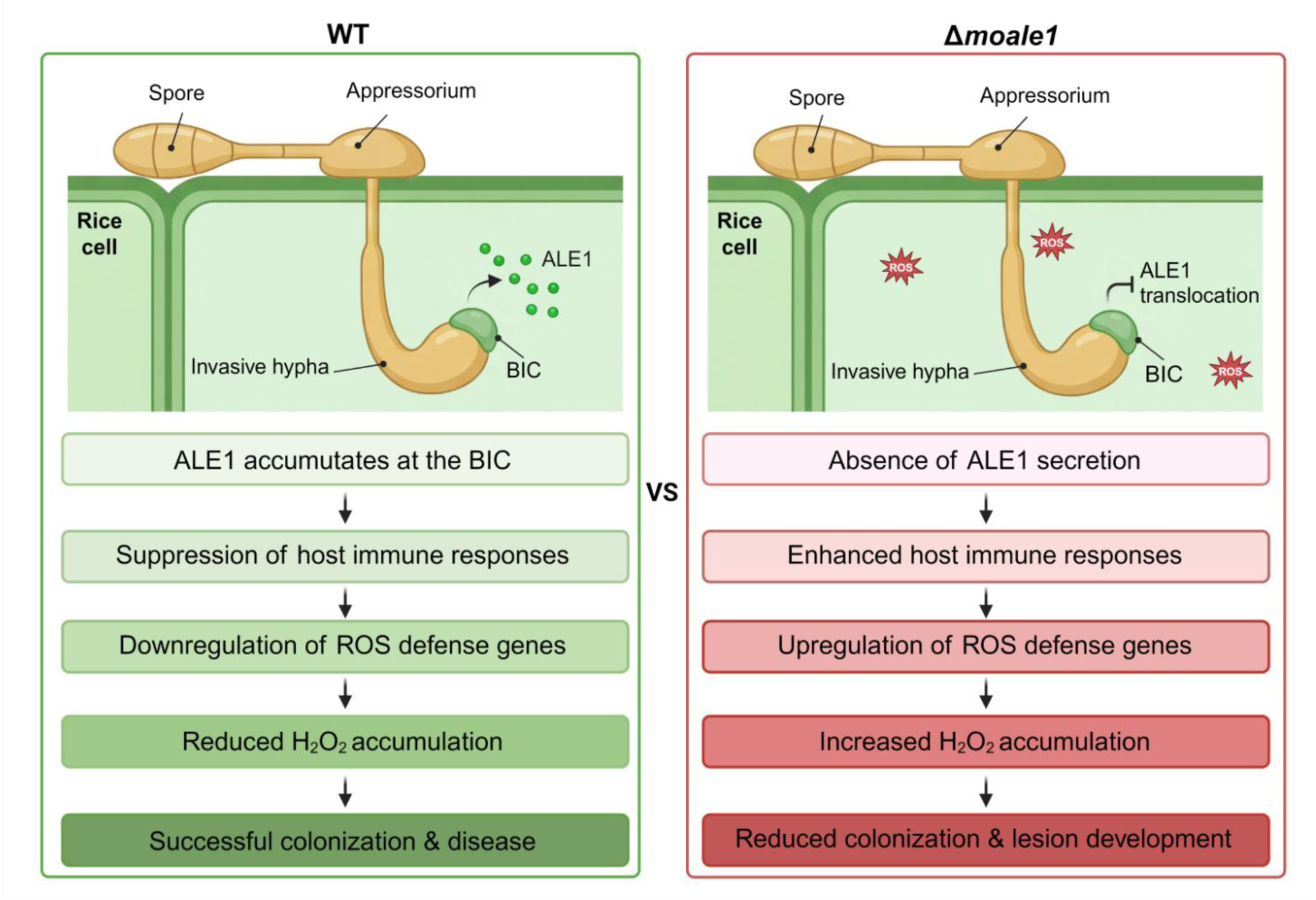
Proposed model for ALE1 function during rice infection. ALE1 accumulates at the BIC and suppresses host immune responses, reducing ROS-associated defenses and promoting fungal colonization. In the absence of ALE1 (Δ*ale1*), increased expression of ROS-associated defense genes and H₂O₂ accumulation enhance host immunity, leading to reduced lesion development and virulence.

Manipulation of ROS production is a common strategy employed by *M. oryzae* effectors to facilitate infection (Hu et al., 2022; Liu et al., 2022; Shi et al., 2018; Song et al., 2024; Yang et al., 2019). Interestingly, despite elevated H₂O₂ accumulation in Δ*moale1*-infected tissues, we observed no defects in penetration efficiency, invasive hyphal growth, or early cell-to-cell spread. This suggests that the elevated H₂O₂ detected at 30 hpi does not directly restrict fungal growth during the early stages of infection. Instead, host-derived H₂O₂ may function as an immune signal that amplifies downstream defense responses. In plants, ROS signaling promotes numerous immune outputs, including transcriptional reprogramming, callose deposition, and programmed cell death (Kou et al., 2019; Qiu et al., 2019; Wang et al., 2019). Consequently, the increased H₂O₂ accumulation observed in Δ*moale1*-infected tissues may indirectly contribute to disease resistance by activating additional defense pathways that become effective later in the infection process. Future studies examining fungal biomass accumulation, callose deposition, and other defense outputs throughout infection will help clarify how MoALE1-mediated suppression of ROS contributes to pathogenicity.

Similar suppression of host immunity has been reported for several characterized *M. oryzae* effectors. Cytoplasmic effectors such as AvrPiz-t interfere with host immune signaling and ROS-associated defenses, whereas nucleus-targeted effectors, including MoHTR1 and MoHTR2, function as transcriptional regulators that alter host gene expression to favor infection (Kim et al., 2020; Park et al., 2012). The observation that MoALE1 suppresses both BAX-mediated cell death and host H₂O₂ accumulation suggests that it likewise functions as an intracellular immune suppressor, although its molecular targets remain unknown.

In the context of these findings, it is important to note that plant pathogen effectors with ADP-ribosyltransferase activity are not uncommon. For instance, in *P. syringae* pv. *tomato*, HopU1 exhibits mono-ADP-ribosyltransferase activity, modifying the host glycine-rich RNA-binding protein GRP7 at R49. This modification interferes with GRP7-mediated regulation of immune receptor transcripts, ultimately impairing host recognition of the pathogen (Jeong et al., 2011; Nicaise et al., 2013). Additional ART effectors have also been identified in *P. syringae*, including AvrRpm1, which targets the callose deposition regulator AtRIN4, and HOF2, which ADP-ribosylates the host kinase MKK5, a key regulator of immune signaling (Redditt et al., 2019; Wang et al., 2010). Similarly, in *S. scabies*, Scabin possesses an ARTC domain and catalyzes ADP-ribosylation of guanosine-containing substrates, including DNA, RNA, GDP, and cGMP (Lyons et al., 2016). These examples highlight the diverse roles of ADP-ribosylation in microbial virulence. However, the majority of characterized ART effectors are confined to bacterial pathogens, leaving experimentally validated fungal effectors with ADP-ribosyltransferase activity largely absent from the literature.

Although MoALE1 contains an ART-like fold and shares structural similarity with members of the heat-labile enterotoxin/cholera toxin family, we were unable to experimentally validate NAD⁺ binding or catalytic activity (Lüscher et al., 2022; Wu et al., 2024). One possible explanation is that MoALE1 requires a host-derived substrate, cofactor, or interacting partner that was absent from the biochemical assays performed in this study. Therefore, the identification and experimental validation of MoALE1-interacting proteins represent an important future direction. Such studies may reveal whether MoALE1 functions as a bona fide ADP-ribosyltransferase or whether its ART-like fold has evolved an alternative role in fungal pathogenesis. MoALE1 also represents one of the first experimentally characterized effectors predicted to possess an ART-like fold in *M. oryzae*. Previous analyses identified between 10 and 27 candidate effectors containing predicted ADP-ribosyltransferase-like folds within the *M. oryzae* genome (Seong & Krasileva, 2021). Whether these proteins share common functions, target similar host pathways, or exhibit functional redundancy remains unknown. The observation that putative ART effectors, like MoALE1 and Mep2, localize to the BIC during biotrophy raises the possibility that ART-like effectors comprise a broader class of intracellular virulence factors in *M. oryzae* (Yan et al., 2023). Future characterization of additional ART-like effectors will be necessary to determine the evolutionary and functional significance of this emerging effector group.

In summary, our study identifies MoALE1 as a biotrophy-associated cytoplasmic effector that contributes to rice blast pathogenicity by suppressing host immune responses (Fig. 7). MoALE1 is expressed during biotrophy, accumulates at the BIC, suppresses BAX-mediated cell death, and limits host H₂O₂ accumulation during infection. Although its biochemical activity remains unknown, the predicted ART-like fold and its contribution to virulence make MoALE1 an attractive target for future investigation. Continued characterization of MoALE1 and related ART-like effectors will provide new insight into the molecular mechanisms underlying *M. oryzae* pathogenesis and may reveal novel targets for rice blast disease management.

## CRediT authorship contribution statement

**Jessie Fernandez**: Conceptualization, Methodology, Formal analysis, Validation, Resources, Funding acquisition, Project administration, Supervision, Writing – original draft, Writing – review & editing. **Rachel E. Kalicharan**: Methodology, Investigation, Data curation, Writing – original draft, Writing – review & editing. **Nalleli Payne**: Investigation, Writing – review & editing. **Isabela G. Tagala**: Investigation, Writing – review & editing. **Krzysztof Pawlowski**: Formal analysis, Writing – review & editing.

## Acknowledgements

We thank the members of the Fernandez Laboratory for their valuable discussions, constructive feedback, and comments on earlier versions of the manuscript. All authors reviewed and approved the final manuscript.

## Declaration of Competing interest

The authors declare no competing interests.

## Funding

This work was supported by the University of Florida Office of Research Opportunity Seed Fund (ROSF) [2022-00131629], the University of Florida Institute of Food and Agricultural Sciences (UF/IFAS) Early Career Seed Grant Program, and the Research Capacity Fund (Hatch) Program (FLA-MCS-006136) from the U.S. Department of Agriculture National Institute of Food and Agriculture (USDA-NIFA).

## Data availability

Data will be made available on request.

Appendix A. Supplementary data

## Notes

### Competing Interest Statement

The authors have declared no competing interest.

## References

Abramson, J., Adler, J., Dunger, J., Evans, R., Green, T., Pritzel, A., Ronneberger, O., Willmore, L., Ballard, A. J., Bambrick, J., Bodenstein, S. W., Evans, D. A., Hung, C.-C., O’Neill, M., Reiman, D., Tunyasuvunakool, K., Wu, Z., Žemgulytė, A., Arvaniti, E., Beattie, C., Bertolli, O., Bridgland, A., Cherepanov, A., Congreve, M., Cowen-Rivers, A. I., Cowie, A., Figurnov, M., Fuchs, F. B., Gladman, H., Jain, R., Khan, Y. A., Low, C. M. R., Perlin, K., Potapenko, A., Savy, P., Singh, S., Stecula, A., Thillaisundaram, A., Tong, C., Yakneen, S., Zhong, E. D., Zielinski, M., Žídek, A., Bapst, V., Kohli, P., Jaderberg, M., Hassabis, D., & Jumper, J. M. 2024. Accurate structure prediction of biomolecular interactions with AlphaFold 3. Nature, 630, 493–500. 10.1038/s41586-024-07487-w

Almeida, F., Rodrigues, M. L., & Coelho, C. 2019. The still underestimated problem of fungal diseases worldwide. Frontiers in Microbiology, 10, 214. 10.3389/fmicb.2019.00214

Aravind, L., Zhang, D., De Souza, R. F., Anand, S., & Iyer, L. M. 2014. The natural history of ADP-ribosyltransferases and the ADP-ribosylation system. In F. Koch-Nolte (Ed.), Endogenous ADP-Ribosylation (Vol. 384, pp. 3–32). Springer International Publishing. 10.1007/82_2014_414

Black, M. H., Osinski, A., Park, G. J., Gradowski, M., Servage, K. A., Pawłowski, K., & Tagliabracci, V. S. 2021. A *Legionella* effector ADP-ribosyltransferase inactivates glutamate dehydrogenase. J Biol Chem, 296, 100301. 10.1016/j.jbc.2021.100301

Chappidi, N., Quail, T., Doll, S., Vogel, L. T., Aleksandrov, R., Felekyan, S., Kühnemuth, R., Stoynov, S., Seidel, C. A. M., Brugués, J., Jahnel, M., Franzmann, T. M., & Alberti, S. 2024. PARP1-DNA co-condensation drives DNA repair site assembly to prevent disjunction of broken DNA ends. Cell, 187, 945–961.e918. 10.1016/j.cell.2024.01.015

Chen, X., Laborda, P., Dong, Y., & Liu, F. 2020. Evaluation of suitable reference genes for normalization of quantitative real-time PCR analysis in rice plants under *Xanthomonas oryzae* pv. oryzae-infection and melatonin supplementation. *Food Production*, Processing and Nutrition, 2, 21. 10.1186/s43014-020-00035-9

Dasovich, M., & Leung, A. K. L. 2023. PARPs and ADP-ribosylation: Deciphering the complexity with molecular tools. Molecular Cell, 83, 1552–1572. 10.1016/j.molcel.2023.04.009

De Jong, J. C., McCormack, B. J., Smirnoff, N., & Talbot, N. J. 1997. Glycerol generates turgor in rice blast. Nature, 389, 244–244. 10.1038/38418

Eseola, A. B., Ryder, L. S., Osés-Ruiz, M., Findlay, K., Yan, X., Cruz-Mireles, N., Molinari, C., Garduño-Rosales, M., & Talbot, N. J. 2021. Investigating the cell and developmental biology of plant infection by the rice blast fungus *Magnaporthe oryzae*. Fungal Genetics and Biology, 154, 103562. 10.1016/j.fgb.2021.103562

Fernandez, J., & Orth, K. 2018. Rise of a cereal killer: The biology of *Magnaporthe oryzae* biotrophic growth. Trends in Microbiology, 26, 582–597. 10.1016/j.tim.2017.12.007

Garcia, N., Farmer, A., Baptiste, R., & Fernandez, J. 2023. Gene Replacement by a Selectable Marker in the Filamentous Fungus Magnaporthe oryzae. BIO-PROTOCOL, 13. 10.21769/BioProtoc.4809

Giraldo, M. C., Dagdas, Y. F., Gupta, Y. K., Mentlak, T. A., Yi, M., Martinez-Rocha, A. L., Saitoh, H., Terauchi, R., Talbot, N. J., & Valent, B. 2013. Two distinct secretion systems facilitate tissue invasion by the rice blast fungus *Magnaporthe oryzae*. Nature Communications, 4, 1996. 10.1038/ncomms2996

Godfray, H. C. J., Mason-D’Croz, D., & Robinson, S. 2016. Food system consequences of a fungal disease epidemic in a major crop. Philosophical Transactions of the Royal Society B: Biological Sciences, 371, 20150467. 10.1098/rstb.2015.0467

Goodin, M. M., Dietzgen, R. G., Schichnes, D., Ruzin, S., & Jackson, A. O. 2002. pGD vectors: versatile tools for the expression of green and red fluorescent protein fusions in agroinfiltrated plant leaves. Plant J, 31, 375–383. 10.1046/j.1365-313x.2002.01360.x

Hasnat, S., Hoque, M. N., Mahbub, M. M., Jummah, J. B., Ali, J., Sakif, T. I., Crandal, K. A., & Islam, T. 2025. In-silico identification and characterization of effector proteins in the rice blast pathogen *Magnaporthe oryzae*. Computational and Structural Biotechnology Reports, 2, 100028. 10.1016/j.csbr.2024.100028

Hu, J., Liu, M., Zhang, A., Dai, Y., Chen, W., Chen, F., Wang, W., Shen, D., Telebanco-Yanoria, M. J., Ren, B., Zhang, H., Zhou, H., Zhou, B., Wang, P., & Zhang, Z. 2022. Co-evolved plant and blast fungus ascorbate oxidases orchestrate the redox state of host apoplast to modulate rice immunity. Molecular Plant, 15, 1347–1366. 10.1016/j.molp.2022.07.001

Jeong, B.-r., Lin, Y., Joe, A., Guo, M., Korneli, C., Yang, H., Wang, P., Yu, M., Cerny, R. L., Staiger, D., Alfano, J. R., & Xu, Y. 2011. Structure function analysis of an ADP-ribosyltransferase type III effector and its RNA-binding target in plant immunity. Journal of Biological Chemistry, 286, 43272–43281. 10.1074/jbc.M111.290122

Jørgensen, R., Purdy, A. E., Fieldhouse, R. J., Kimber, M. S., Bartlett, D. H., & Merrill, A. R. 2008. Cholix Toxin, a novel ADP-ribosylating factor from *Vibrio cholerae*. Journal of Biological Chemistry, 283, 10671–10678. 10.1074/jbc.M710008200

Kankanala, P., Czymmek, K., & Valent, B. 2007. Roles for rice membrane dynamics and plasmodesmata during biotrophic invasion by the blast fungus. Plant Cell, 19, 706–724. 10.1105/tpc.106.046300

Kershaw, M. J., & Talbot, N. J. 2009. Genome-wide functional analysis reveals that infection-associated fungal autophagy is necessary for rice blast disease. Proceedings of the National Academy of Sciences, 106, 15967–15972. 10.1073/pnas.0901477106

Khang, C. H., Berruyer, R., Giraldo, M. C., Kankanala, P., Park, S.-Y., Czymmek, K., Kang, S., & Valent, B. 2010. Translocation of *Magnaporthe oryzae* effectors into rice cells and their subsequent cell-to-cell movement. The Plant Cell, 22, 1388–1403. 10.1105/tpc.109.069666

Khaskhali, S., Xiao, X., Zhang, Z., Solangi, F., Hussain, S., & Chen, Y. 2025. Expression profile and characterization of respiratory burst oxidase homolog genes in rice under MeJA, SA and Xoo treatments. Scientific Reports, 15, 5936. 10.1038/s41598-025-88731-9

Kim, S., Kim, C.-Y., Park, S.-Y., Kim, K.-T., Jeon, J., Chung, H., Choi, G., Kwon, S., Choi, J., Jeon, J., Jeon, J.-S., Khang, C. H., Kang, S., & Lee, Y.-H. 2020. Two nuclear effectors of the rice blast fungus modulate host immunity via transcriptional reprogramming. Nature Communications, 11, 5845. 10.1038/s41467-020-19624-w

Kou, Y., Qiu, J., & Tao, Z. 2019. Every Coin Has Two Sides: Reactive Oxygen Species during rice-*Magnaporthe oryzae* Interaction. International Journal of Molecular Sciences, 20, 1191. 10.3390/ijms20051191

Lee, S.-J., & Rose, J. K. C. 2012. A yeast secretion trap assay for identification of secreted proteins from eukaryotic phytopathogens and their plant hosts. In M. D. Bolton & B. P. H. J. Thomma (Eds.), Plant Fungal Pathogens (Vol. 835, pp. 519–530). Humana Press. 10.1007/978-1-61779-501-5_32

Lee, S., Völz, R., Lim, Y. J., Harris, W., Kim, S., & Lee, Y. H. 2023. The nuclear effector MoHTR3 of *Magnaporthe oryzae* modulates host defence signalling in the biotrophic stage of rice infection. Molecular Plant Pathology, 24, 602–615. 10.1111/mpp.13326

Liu, X., Gao, Y., Guo, Z., Wang, N., Wegner, A., Wang, J., Zou, X., Hu, J., Liu, M., Zhang, H., Zheng, X., Wang, P., Schaffrath, U., & Zhang, Z. 2022. MoIug4 is a novel secreted effector promoting rice blast by counteracting host OsAHL1-regulated ethylene gene transcription. New Phytologist, 235, 1163–1178. 10.1111/nph.18169

Lüscher, B., Ahel, I., Altmeyer, M., Ashworth, A., Bai, P., Chang, P., Cohen, M., Corda, D., Dantzer, F., Daugherty, M. D., Dawson, T. M., Dawson, V. L., Deindl, S., Fehr, A. R., Feijs, K. L. H., Filippov, D. V., Gagné, J. P., Grimaldi, G., Guettler, S., Hoch, N. C., Hottiger, M. O., Korn, P., Kraus, W. L., Ladurner, A., Lehtiö, L., Leung, A. K. L., Lord, C. J., Mangerich, A., Matic, I., Matthews, J., Moldovan, G. L., Moss, J., Natoli, G., Nielsen, M. L., Niepel, M., Nolte, F., Pascal, J., Paschal, B. M., Pawłowski, K., Poirier, G. G., Smith, S., Timinszky, G., Wang, Z. Q., Yélamos, J., Yu, X., Zaja, R., & Ziegler, M. 2022. ADP-ribosyltransferases, an update on function and nomenclature. The FEBS Journal, 289, 7399–7410. 10.1111/febs.16142

Lüscher, B., Bütepage, M., Eckei, L., Krieg, S., Verheugd, P., & Shilton, B. H. 2018. ADP-Ribosylation, a multifaceted posttranslational modification involved in the control of cell physiology in health and disease. Chemical Reviews, 118, 1092–1136. 10.1021/acs.chemrev.7b00122

Lyons, B., Ravulapalli, R., Lanoue, J., Lugo, M. R., Dutta, D., Carlin, S., & Merrill, A. R. 2016. Scabin, a Novel DNA-acting ADP-ribosyltransferase from *Streptomyces scabies*. J Biol Chem, 291, 11198–11215. 10.1074/jbc.M115.707653

Meng, E. C., Goddard, T. D., Pettersen, E. F., Couch, G. S., Pearson, Z. J., Morris, J. H., & Ferrin, T. E. 2023. UCSF ChimeraX: Tools for structure building and analysis. Protein Science, 32, e4792. 10.1002/pro.4792

Mentlak, T. A., Kombrink, A., Shinya, T., Ryder, L. S., Otomo, I., Saitoh, H., Terauchi, R., Nishizawa, Y., Shibuya, N., Thomma, B. P. H. J., & Talbot, N. J. 2012. Effector-mediated suppression of chitin-triggered immunity by *Magnaporthe oryzae* Is necessary for rice blast disease. The Plant Cell, 24, 322–335. 10.1105/tpc.111.092957

Mikolčević, P., Hloušek-Kasun, A., Ahel, I., & Mikoč, A. 2021. ADP-ribosylation systems in bacteria and viruses. Computational and Structural Biotechnology Journal, 19, 2366–2383. 10.1016/j.csbj.2021.04.023

Mosquera, G., Giraldo, M. C., Khang, C. H., Coughlan, S., & Valent, B. 2009. Interaction transcriptome analysis identifies *Magnaporthe oryzae* BAS1-4 as biotrophy-associated secreted proteins in rice blast disease. The Plant Cell, 21, 1273–1290. 10.1105/tpc.107.055228

Nicaise, V., Joe, A., Jeong, B.-r., Korneli, C., Boutrot, F., Westedt, I., Staiger, D., Alfano, J. R., & Zipfel, C. 2013. *Pseudomonas* HopU1 modulates plant immune receptor levels by blocking the interaction of their mRNAs with GRP7. The EMBO Journal, 32, 701–712. 10.1038/emboj.2013.15

Omar, S. C., Bentley, M. A., Morieri, G., Preston, G. M., & Gurr, S. J. 2016. Validation of reference genes for robust qRT-PCR gene expression analysis in the rice blast fungus *Magnaporthe oryzae*. PLOS ONE, 11, e0160637. 10.1371/journal.pone.0160637

Osés-Ruiz, M., Sakulkoo, W., Littlejohn, G. R., Martin-Urdiroz, M., & Talbot, N. J. 2017. Two independent S-phase checkpoints regulate appressorium-mediated plant infection by the rice blast fungus *Magnaporthe oryzae*. Proceedings of the National Academy of Sciences, 114. 10.1073/pnas.1611307114

Park, C.-H., Chen, S., Shirsekar, G., Zhou, B., Khang, C. H., Songkumarn, P., Afzal, A. J., Ning, Y., Wang, R., Bellizzi, M., Valent, B., & Wang, G.-L. 2012. The *Magnaporthe oryzae* effector AvrPiz-t targets the RING E3 Ubiquitin Ligase APIP6 to suppress pathogen-associated molecular pattern–triggered immunity in rice. The Plant Cell, 24, 4748–4762. 10.1105/tpc.112.105429

Park, C. H., Shirsekar, G., Bellizzi, M., Chen, S., Songkumarn, P., Xie, X., Shi, X., Ning, Y., Zhou, B., Suttiviriya, P., Wang, M., Umemura, K., & Wang, G.-L 2016. The E3 ligase APIP10 connects the effector AvrPiz-t to the NLR receptor Piz-t in rice. PLOS Pathogens, 12, e1005529. 10.1371/journal.ppat.1005529

Peng, X., Hu, Y., Tang, X., Zhou, P., Deng, X., Wang, H., & Guo, Z. 2012. Constitutive expression of rice WRKY30 gene increases the endogenous jasmonic acid accumulation, PR gene expression and resistance to fungal pathogens in rice. Planta, 236, 1485–1498. 10.1007/s00425-012-1698-7

Qiu, Z., Zhu, L., He, L., Chen, D., Zeng, D., Chen, G., Hu, J., Zhang, G., Ren, D., Dong, G., Gao, Z., Shen, L., Zhang, Q., Guo, L., & Qian, Q. 2019. DNA damage and reactive oxygen species cause cell death in the rice local lesions 1 mutant under high light and high temperature. New Phytologist, 222, 349–365. 10.1111/nph.15597

Redditt, T. J., Chung, E.-H., Zand Karimi, H., Rodibaugh, N., Zhang, Y., Trinidad, J. C., Kim, J. H., Zhou, Q., Shen, M., Dangl, J. L., Mackey, D. M., & Innes, R. W. 2019. AvrRpm1 functions as an ADP-ribosyl transferase to modify NOI domain-containing proteins, including *Arabidopsis* and soybean RPM1-interacting protein 4. The Plant Celltpc.00020.02019. 10.1105/tpc.19.00020

Robert, X., Guillon, C., & Gouet, P. 2025. FoldScript: a web server for the efficient analysis of AI-generated 3D protein models. Nucleic Acids Research, 53, W277–W282. 10.1093/nar/gkaf326

Rosado, M. M., & Pioli, C. 2021. ADP-ribosylation in evasion, promotion and exacerbation of immune responses. Immunology, 164, 15–30. 10.1111/imm.13332

Sakulkoo, W., Osés-Ruiz, M., Oliveira Garcia, E., Soanes, D. M., Littlejohn, G. R., Hacker, C., Correia, A., Valent, B., & Talbot, N. J. 2018. A single fungal MAP kinase controls plant cell-to-cell invasion by the rice blast fungus. Science, 359, 1399–1403. 10.1126/science.aaq0892

Schindelin, J., Arganda-Carreras, I., Frise, E., Kaynig, V., Longair, M., Pietzsch, T., Preibisch, S., Rueden, C., Saalfeld, S., Schmid, B., Tinevez, J.-Y., White, D. J., Hartenstein, V., Eliceiri, K., Tomancak, P., & Cardona, A. 2012. Fiji: an open-source platform for biological-image analysis. Nature Methods, 9, 676–682. 10.1038/nmeth.2019

Seong, K., & Krasileva, K. V. 2021. Computational structural genomics unravels common folds and novel families in the secretome of fungal phytopathogen. Mol Plant Microbe Interact, 34, 1267–1280. 10.1094/MPMI-03-21-0071-R

Seong, K., & Krasileva, K. V. 2023. Prediction of effector protein structures from fungal phytopathogens enables evolutionary analyses. Nature Microbiology, 8, 174–187. 10.1038/s41564-022-01287-6

Sharpee, W., Oh, Y., Yi, M., Franck, W., Eyre, A., Okagaki, L. H., Valent, B., & Dean, R. A. 2017. Identification and characterization of suppressors of plant cell death (SPD) effectors from *Magnaporthe oryzae*. Molecular Plant Pathology, 18, 850–863. 10.1111/mpp.12449

Shi, X., Long, Y., He, F., Zhang, C., Wang, R., Zhang, T., Wu, W., Hao, Z., Wang, Y., Wang, G.-L., & Ning, Y. 2018. The fungal pathogen *Magnaporthe oryzae* suppresses innate immunity by modulating a host potassium channel. PLOS Pathogens, 14, e1006878. 10.1371/journal.ppat.1006878

Sievers, F., Wilm, A., Dineen, D., Gibson, T. J., Karplus, K., Li, W., Lopez, R., McWilliam, H., Remmert, M., Söding, J., Thompson, J. D., & Higgins, D. G. 2011. Fast, scalable generation of high-quality protein multiple sequence alignments using Clustal Omega. Molecular Systems Biology, 7, MSB201175. 10.1038/msb.2011.75

Song, L., Yang, T., Wang, X., Ye, W., & Lu, G. 2024. *Magnaporthe oryzae* Effector AvrPik-D Targets Rice Rubisco Small Subunit OsRBCS4 to Suppress Immunity. Plants, 13, 1214. 10.3390/plants13091214

Spence, C., Alff, E., Johnson, C., Ramos, C., Donofrio, N., Sundaresan, V., & Bais, H. 2014. Natural rice rhizospheric microbes suppress rice blast infections. BMC Plant Biology, 14, 130. 10.1186/1471-2229-14-130

Teufel, F., Armenteros, J. J. A., Johansen, A. R., Gíslason, M. H., Pihl, S. I., Tsirigos, K. D., Winther, O., Brunak, S., Heijne, G. V., & Nielsen, H. 2022. SignalP 6.0 predicts all five types of signal peptides using protein language models. Nature Biotechnology, 40, 1023–1025. 10.1038/s41587-021-01156-3

Veneault-Fourrey, C., Barooah, M., Egan, M., Wakley, G., & Talbot, N. J. 2006. Autophagic fungal cell death is necessary for infection by the rice blast fungus. Science, 312, 580–583. 10.1126/science.1124550

Wang, C., Liu, Y., Liu, L., Wang, Y., Yan, J., Wang, C., Li, C., & Yang, J. 2019. The biotrophy-associated secreted protein 4 (BAS4) participates in the transition of *Magnaporthe oryzae* from the biotrophic to the necrotrophic phase. Saudi Journal of Biological Sciences, 26, 795–807. 10.1016/j.sjbs.2019.01.003

Wang, Y., Li, J., Hou, S., Wang, X., Li, Y., Ren, D., Chen, S., Tang, X., & Zhou, J.-M. 2010. A *Pseudomonas syringae* ADP-ribosyltransferase inhibits *Arabidopsis* mitogen-activated protein kinase kinases. The Plant Cell, 22, 2033–2044. 10.1105/tpc.110.075697

Wilson, R. A., & Talbot, N. J. 2009. Under pressure: investigating the biology of plant infection by *Magnaporthe oryzae*. Nature Reviews Microbiology, 7, 185–195. 10.1038/nrmicro2032

Wu, H., Lu, A., Yuan, J., Yu, Y., Lv, C., & Lu, J. 2024. Mono-ADP-ribosylation, a MARylationmultifaced modification of protein, DNA and RNA: characterizations, functions and mechanisms. Cell Death Discovery, 10, 226. 10.1038/s41420-024-01994-5

Xu, D., Jaroszewski, L., Li, Z., & Godzik, A. 2014. FFAS-3D: improving fold recognition by including optimized structural features and template re-ranking. Bioinformatics, 30, 660–667. 10.1093/bioinformatics/btt578

Yan, X., Tang, B., Ryder, L. S., MacLean, D., Were, V. M., Eseola, A. B., Cruz-Mireles, N., Ma, W., Foster, A. J., Osés-Ruiz, M., & Talbot, N. J. 2023. The transcriptional landscape of plant infection by the rice blast fungus *Magnaporthe oryzae* reveals distinct families of temporally co-regulated and structurally conserved effectors. The Plant Cell, 35, 1360–1385. 10.1093/plcell/koad036

Yang, C., Li, W., Cao, J., Meng, F., Yu, Y., Huang, J., Jiang, L., Liu, M., Zhang, Z., Chen, X., Miyamoto, K., Yamane, H., Zhang, J., Chen, S., & Liu, J. 2017. Activation of ethylene signaling pathways enhances disease resistance by regulating ROS and phytoalexin production in rice. The Plant Journal, 89, 338–353. 10.1111/tpj.13388

Yang, C., Yu, Y., Huang, J., Meng, F., Pang, J., Zhao, Q., Islam, M. A., Xu, N., Tian, Y., & Liu, J. 2019. Binding of the *Magnaporthe oryzae* chitinase MoChia1 by a rice tetratricopeptide repeat protein allows free chitin to trigger immune responses. The Plant Cell, 31, 172–188. 10.1105/tpc.18.00382

Zhang, H.-f., Islam, T., & Liu, W.-d. 2022. Integrated pest management programme for cereal blast fungus *Magnaporthe oryzae*. Journal of Integrative Agriculture, 21, 3420–3433. 10.1016/j.jia.2022.08.056

Zhong, Z., Chen, M., Lin, L., Han, Y., Bao, J., Tang, W., Lin, L., Lin, Y., Somai, R., Lu, L., Zhang, W., Chen, J., Hong, Y., Chen, X., Wang, B., Shen, W.-C., Lu, G., Norvienyeku, J., Ebbole, D. J., & Wang, Z. 2018. Population genomic analysis of the rice blast fungus reveals specific events associated with expansion of three main clades. The ISME Journal, 12, 1867–1878. 10.1038/s41396-018-0100-6

Zimmermann, L., Stephens, A., Nam, S. Z., Rau, D., Kübler, J., Lozajic, M., Gabler, F., Söding, J., Lupas, A. N., & Alva, V. 2018. A Completely Reimplemented MPI Bioinformatics Toolkit with a New HHpred Server at its Core. J Mol Biol, 430, 2237–2243. 10.1016/j.jmb.2017.12.007

