## Supplemental figures for "MoALE1, an ART-like effector, contributes to rice blast virulence through host immune suppression"

### Supplemental Materials

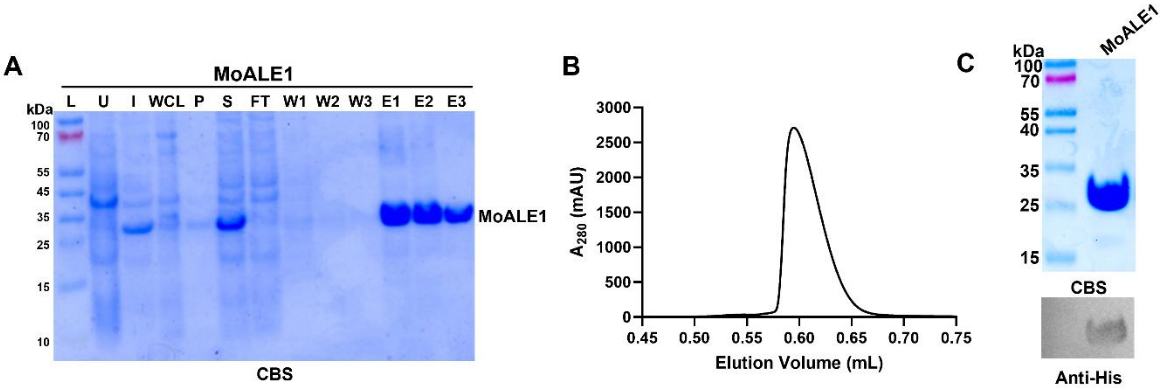


**Figure S1.** Expression and purification of recombinant MoALE1. **(A)** Coomassie Brilliant Blue-stained (CBS) SDS-PAGE analysis of recombinant His-tagged MoALE1 purified by Ni-NTA affinity chromatography. L, molecular weight marker; U, uninduced; I, induced; WCL, whole-cell lysate; P, pellet; S, soluble fraction; FT, flow-through; W1-W3, wash fractions; E1-E3, elution fractions. **(B)** Size-exclusion chromatography profile of purified MoALE1 monitored at 280 nm. **(C)** CBS-stained SDS-PAGE gel (top) and anti-His immunoblot (bottom) confirming purification and identity of recombinant MoALE1 protein, which migrated at the expected molecular weight (~29 kDa).

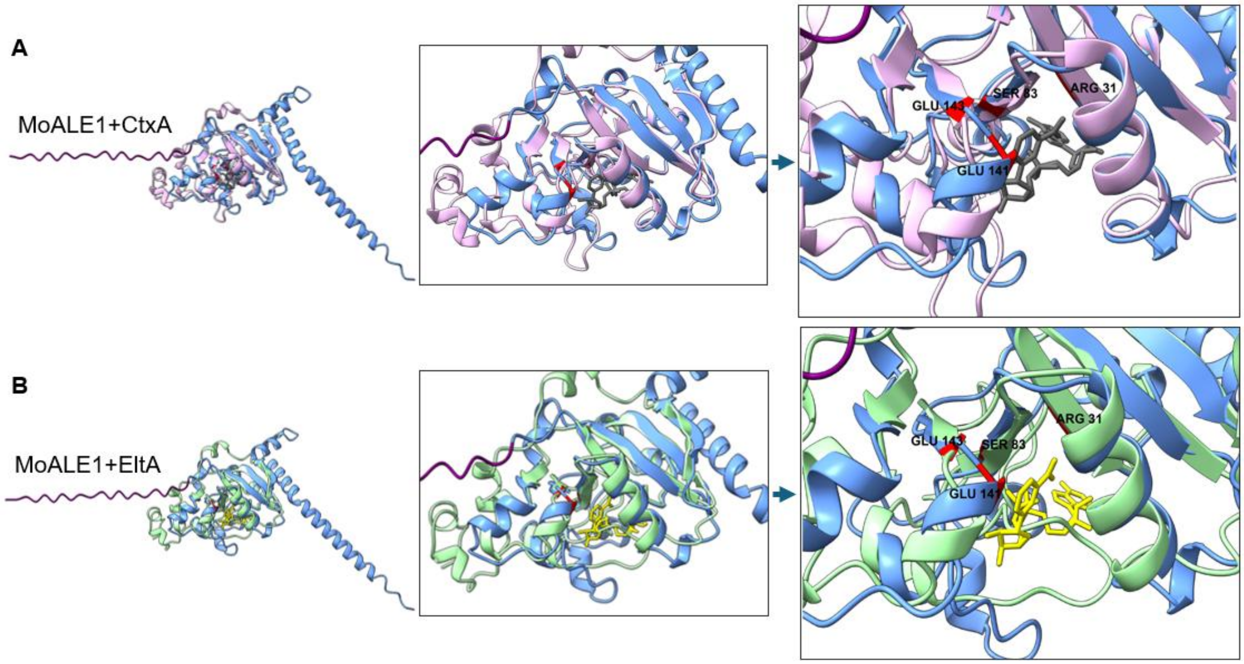


**Figure S2.** Structural similarity of MoALE1 to bacterial enterotoxin ADP-ribosyltransferases. **(A-B)** Structural superposition of the AlphaFold3-predicted MoALE1 model (blue) with the catalytic A subunits of cholera toxin (CtxA, pink) and of heat-labile enterotoxin (EltA, green); RMSD values are 0.891 Å and 1.086 Å, respectively. In (A) binding configuration of $NAD^+$ (gray) is based on the CtxA crystal structure (PDB: 2A5F), in (B) predicted $NAD^+$ is shown in yellow. Conserved residues comprising the predicted R-S-ExE motif (R31, S83, E141, and E143) are highlighted in red and shown in proximity to the predicted $NAD^+$ molecule. The structural overlap supports conservation of an ART-like fold between MoALE1 and bacterial enterotoxins.

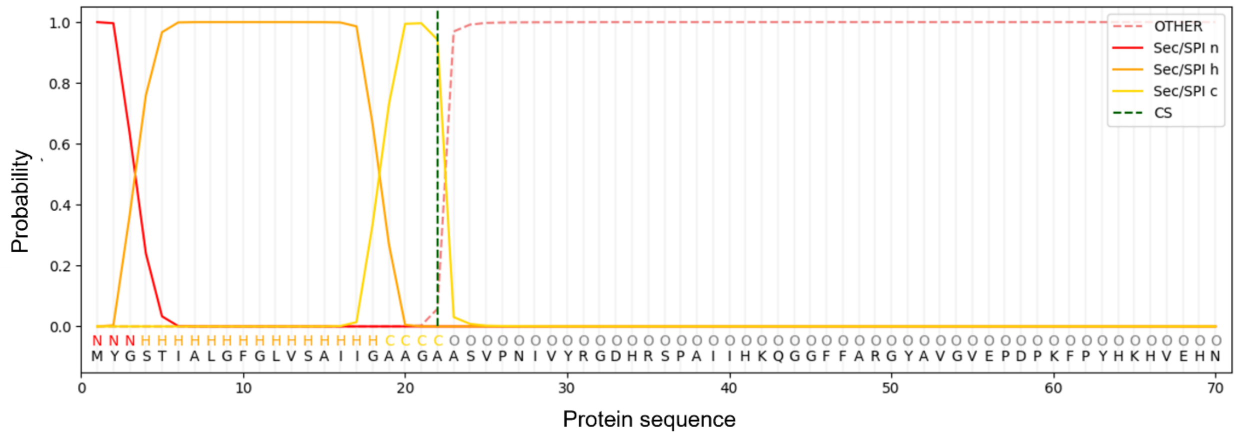


**Figure S3.** Signal peptide prediction for MoALE1. SignalP 6.0 analysis identified an N-terminal secretion signal peptide in MoALE1, with a predicted cleavage site between amino acid residues 22 and 23. The predicted N-, H-, and C-regions of the signal peptide are indicated, supporting secretion of MoALE1 through the classical secretory pathway.

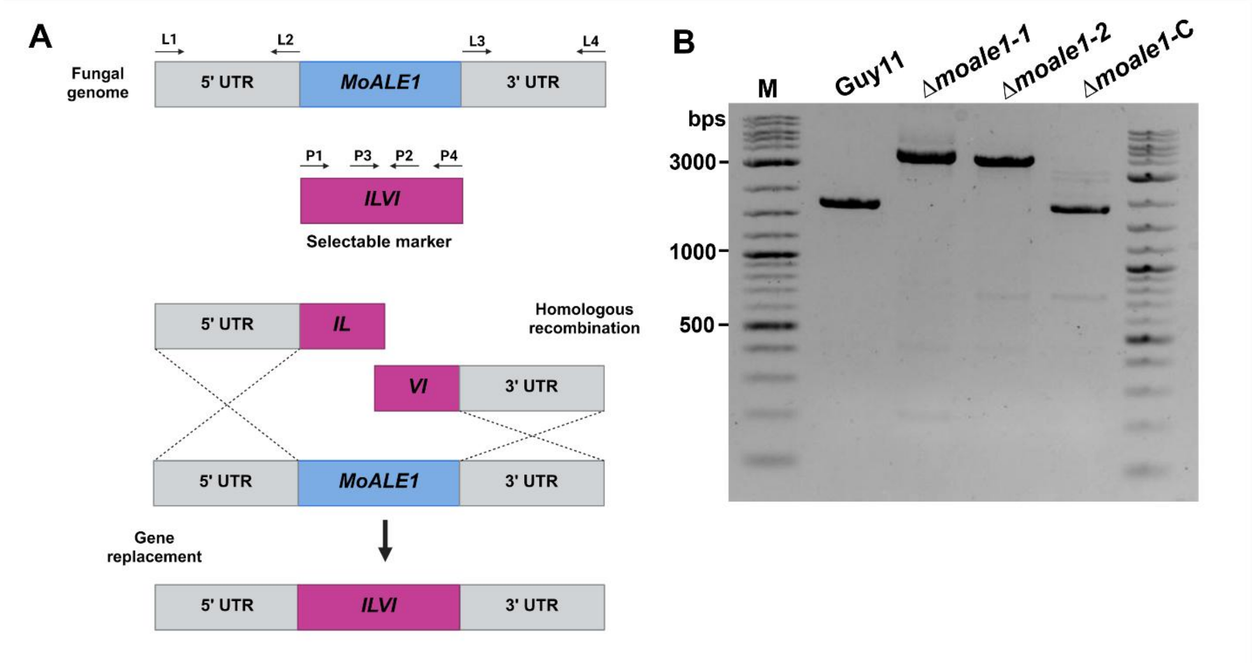


**Figure S4.** Generation and verification of the Δ*moale1* deletion mutant. **(A)** Schematic representation of homologous recombination-mediated replacement of the *MoALE1* locus with the *SUR* selectable marker, which confers sulfonylurea resistance through a mutant *ILV1* allele. Approximately 1-kb regions upstream (5′ UTR) and downstream (3′ UTR) of *MoALE1* were fused to the *SUR* cassette and introduced into *M. oryzae* Guy11 protoplasts. Homologous recombination resulted in replacement of the endogenous *moale1* coding sequence with the *SUR* cassette. Arrows indicate primer positions used in PCR. Created using BioRender.com. **(B)** PCR verification of Δ*moale1* transformants. Amplification of the 5′ and 3′ junction regions confirmed integration of the *SUR* cassette at the *MoALE1* locus, while amplification of the endogenous *moale1* coding sequence confirmed deletion of the gene in mutant strains. M, DNA ladder.

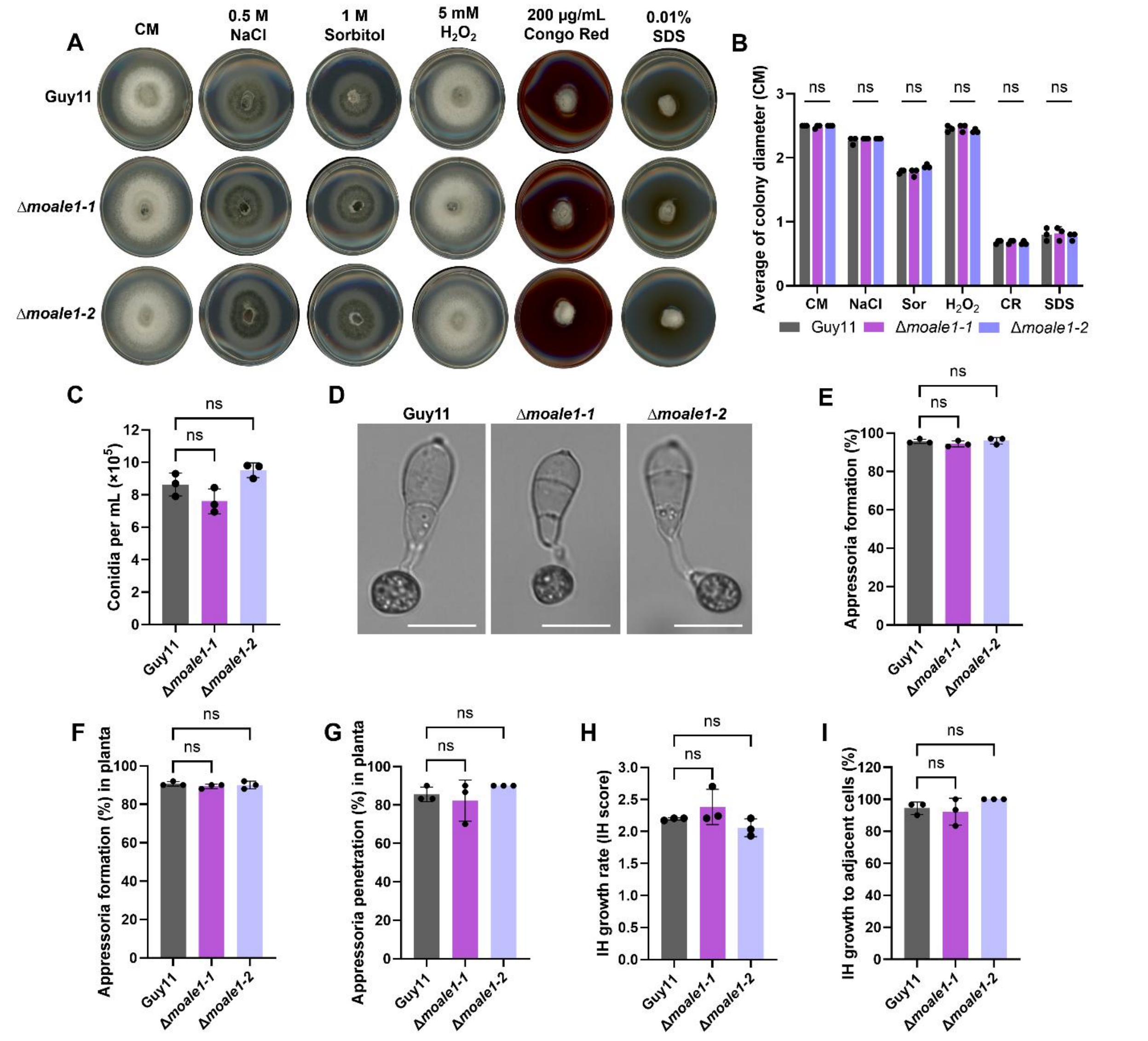


**Figure S5.** *MoALE1* is dispensable for fungal growth, development, and early infection processes. **(A)** *In vitro* stress tolerance assay of Guy11, Δ*moale1*-1, and Δ*moale1*-2 grown on complete medium (CM), osmotic stress medium supplemented with 0.5 M NaCl or 1 M sorbitol, oxidative stress medium supplemented with 5 mM $H_2O_2$, and cell wall stress medium supplemented with 200 μg mL$^{-1}$ Congo Red or 0.01% SDS. Plates were imaged at 5 dpi. **(B)** Quantification of colony diameter under the conditions shown in (A). **(C)** Quantification of sporulation by Guy11, Δ*moale1*-1, and Δ*moale1*-2 following 10 days of growth on CM. **(D)** Representative images of appressoria formed by Guy11, Δ*moale1*-*1*, and Δ*moale1*-2 on hydrophobic coverslips. Scale bar = 10 μm. **(E)** Quantification of *in vitro* appressorium formation on hydrophobic coverslips. **(F)** Quantification of *in planta* appressorium formation. **(G)** Quantification of appressorium penetration efficiency. **(H)** Quantification of invasive hyphal (IH) growth at 24 hpi. IH growth was scored as follows: 0, appressorium formation without penetration; 1, penetration peg formation; 2, primary invasive hypha (PIH); 3, PIH with up to two branches within the primary infected cell; 4,

PIH with three or more branches within the primary infected cell; and 5, spread into adjacent cells. **(I)** Quantification of invasive hyphal spread into adjacent cells at 48 hpi. Each point represents an independent biological replicate (*n* = 3). Bars represent the mean ± SD. Statistical significance was determined using a one-way ANOVA followed by Dunnett's multiple-comparisons test; ns, not significant.
