## Supplemental table S1 for "MoALE1, an ART-like effector, contributes to rice blast virulence through host immune suppression"

**Table S1**. List of primers used in this study.

| Primer Name | Primer Sequences (5’-3’) | Source |
|---|---|---|
| pCB1532-NP-MoALE1-GFP | F: TTACGAGCGGCCGCGGACGTCTGGGAAGATATATTG<br>R: TTACGACTGCAGTAGTAACTGCGGCTGAACATG | This study |
| pYST-MoALE1 | F: CCAAGCTGAATTCAGTCAGCGGCCGCATGTACGGCTCGACAATTG<br>R: GGGTACCTTTTATCCAAGGCGGCCGCTAGTAACTGCGGCTGAAC | This study |
| pYST-MoALE1 NSP | F: CCAAGCTGAATTCAGTCAGCGGCCGCATGGCCAGCGTTCCAAAC<br>R: GGGTACCTTTTATCCAAGGCGGCCGCTAGTAACTGCGGCTGAAC | This study |
| pGD-MoALE1 | F: TAAGCACTCGAGATGTACGGCTCGACAATTGCTTTG<br>R: TAAGCAAAGCTTCTATAGTAACTGCGGCTGAACATGTCG | This study |
| pET28a-MoALE1 | F: GGATCCGAATTCGAGCTCCGGATGGCCAGCGTTCCAAAC<br>R: TGGTGGTGGTGCTCGAGTGCCTATAGTAACTGCGGCTGAAC | This study |
| pET28a-MoALE1E143A | F: CTTTTTCTTACGAAAATGCATTCCTGTCGGATCATCC<br>R: GGATGATCCGACAGGAATGCATTTTCGTAAGAAAAAG | This study |
| MoALE1-LF | F: GTGAGATTTCGTGGCCTTGCC<br>R: GTCGTGACTGGGAAAACCCTGGCGTTGGGATAAAGCCTGTCGGTTTG | This study |
| MoALE1-RF | F: TCCTGTGTGAAATTGTTATCCGCTGTATATTTTGATGTTTGGGGTGGTGGA<br>R: CGATTTCCAAAGCTGTATACCAATCTG | This study |
| MoALE1-Nested | F: AGTGTCGTCGAGAGGGGTGTGAC<br>R: CGATTTCCAAAGCTGTATACCAATCTG | This study |
| qMoALE1 | F: CACTCCAGTCACGCCAGAGAA<br>R: TGAACATGTCGAGCCGCAC | This study |
| qMoACT1 | F: ACAATGGTTCGGGTATGTGC<br>R: CGACAATGGACGGGAAGAC | (Omar et al., 2016) |
| qOsUBQ10 | F: TGGTCAGTAATCAGCCAGTTTGG<br>R: GCACCACAAATACTTGACGAACAG | (Chen et al., 2020) |
| qOsRBOHB | F: TACTTCTACTGGGTGACGAG<br>R: TCTTCACACTAGTTCCCGAC | (Yang et al., 2017) |
| qOsRBOHC | F: CAAGGAGCAAGCAGAGGAATA<br>R: CAGACACAGATGGCAGAAGAA | This study |
| qOsRBOHD | F: GGATTCAACAGGATGTGCAAAG<br>R: CTCTGGTGACCCAGTAGAAATAAG | This study |
| qOsWRKY30 | F: ACTTCTTGAGTCGCCGGTTT<br>R: GCTTCTGGGATGCTCACTGT | (Spence et al., 2014) |
